# Engineering carotenoid and steroidal glycoalkaloid depleted tomato fruit for heterologous production of high value terpenes

**DOI:** 10.64898/2026.05.13.724861

**Authors:** Natalie C. Deans, Jon Cody, Lucas Reist, John P. Hamilton, Colby Starker, Lynn Prichard, Joshua C. Wood, Brieanne Vaillancourt, Björn R. Hamberger, Daniel F. Voytas, C. Robin Buell

**Affiliations:** Center for Applied Genetic Technologies, University of Georgia, Athens, GA, USA; Institute of Plant Breeding, Genetics, and Genomics, University of Georgia, Athens, Georgia, USA; Center for Precision Plant Genomics, Department of Genetics, Cell Biology and Development, University of Minnesota, St. Paul, MN, USA; Department of Biochemistry and Molecular Biology, Molecular Plant Sciences Program, Cell and Molecular Biology Program, Michigan State University, East Lansing, MI, USA; Department of Crop and Soil Sciences, University of Georgia, Athens, GA, USA; The Plant Center, University of Georgia, Athens, Georgia, USA

**Keywords:** Chassis, metabolic engineering, terpene, genome engineering, gene editing, tomato

## Abstract

Plants produce specialized metabolites that function in plant defense and as attractants to pollinators and symbionts. One class of specialized metabolites are terpenoids that are synthesized from universal C_5_ building blocks via activities including terpene synthases, cytochromes P450, and glycosyl transferases. Some terpenes are highly valued for their use as insect repellants, fragrances, antimicrobial compounds, low calorie sweeteners, flavors, and medicines. Low abundance in target tissues, present in complex mixtures, as well as challenging extraction logistics are barriers to economic sustainable production of these compounds from their native species. While heterologous expression of terpenoid biosynthetic genes is feasible, the potential derivation of the products into conjugates via endogenous cytochromes P450 and glycosyl transferases limits this approach. In this project, we used multiplex gene editing technologies to overcome these challenges by creating novel tomato chassis with altered terpenoid biosynthetic capacity in fruit. Excluding central metabolic genes to minimalize impacts on growth and development, we selected 23 known and potential terpene-related genes expressed specifically in the fruit for gene editing. Fruit production and metabolic profiles of three chassis lines with alterations in the major classes of fruit specialized metabolites indicate loss of these genes is tolerated for fruit production. These combinatorial knockouts also showed modulation of native carbon reallocation toward endogenous sinks beneficial for a biosynthetic chassis. Establishing metabolite-modified fruit chassis demonstrates efficient combinatorial editing of entire branches of plant specialized metabolism, facilitating engineering of heterologous terpenes of industrial interest in tomato fruit.

## 1 Introduction

For thousands of years, chemicals derived from plants have been harvested and utilized by humans for their unique natural properties. Beyond nutrition, specialized metabolites (i.e. secondary metabolites) have been used as medicines, dyes, fragrances, insect repellents, and are highly relevant for their industrial uses in modern times. Terpenoids, C_5_-derived molecules, are the most diverse class with ca. 100,000 different structures reported and >76,000 found in plants (DNP, 34.2.; Zhou & Pichersky, 2020a). For example, the sesquiterpenoid viridiflorol, present in the essential oils of several plants and also found in multiple fungal species, is valued as a fragrance, for its antimicrobial properties, and in treatment of multiple disorders including cancer (Akiel et al., 2022; de Matos Balsalobre et al., 2023; Zhang et al., 2021).

In plants, terpenes are synthesized from two parallel precursor pathways operating in the plastid and cytosol in which terpene synthases (TPSs) catalyze cycloisomerization and formation of basic scaffolds (Tholl, 2015). Both pathways form the interconvertible C_5_ isomers isopentenyl diphosphate (IPP) and dimethylallyl diphosphate (DMAPP). In the cytosol, the mevalonate pathway (MVA) synthesizes sesquiterpenes (C_15_) and triterpenes/sterols (C_30_), while the plastidial methylerythritol 4-phosphate (MEP) pathway provides the C_5_ building blocks for mono- (C_10_), di- (C_20_) and tetraterpenes/carotenoids (C_40_) (Zhou & Pichersky, 2020a). These scaffolds are further modified by the action of other enzymes including cytochromes P450 (CYPs/P450s) generating functional groups which can serve as molecular handles for additional conjugation by enzymes such as UTP-dependent glycosyl transferases (UGTs) and acyl transferases (Lanier et al., 2023; J. Wang et al., 2016).

Some terpenes are not cost effective to isolate directly from plants due to high cultivation costs as exemplified by perennials such as white birch, eucalyptus, and plane trees which accumulate betulinic acid used to treat multiple diseases (Aswathy et al., 2022; Lou et al., 2021; Mu et al., 2023). Additionally, biosynthesis at low levels, restriction to specific tissues or cells, or biosynthesis in response to specific stimuli can also prohibit cost effective isolation – as exemplified by the monoterpene indole alkaloids vinblastine and vincristine which are synthesized in rare idioblast cells in *Catharanthus roseus* (Mahroug et al., 2007). Lastly, terpenes can be present in complex mixtures that are challenging or costly to purify.

Heterologous expression platforms in fungi, bacteria, and algae have been used to address some of these challenges. For example, heavy metabolic engineering achieved production of viridiflorol in the bacterial host *E. coli* (Shukal et al., 2019). This required nine enzymes, translational optimization, enzyme engineering (modification of subcellular targeting), inducer optimization, modulated carbon- and co-factor supply, and multi-gene replacement to produce viridiflorol. More complex, further functionalized, terpenoids that are dependent on plant P450-modification remain challenging due to the lack of sufficient reductive equivalents (NADPH and cytochrome P450 reductase, CPR) and their required subcellular compartmentalization (e.g., the endoplasmic reticulum). Additionally, more processing can be necessary for microbial products for the food industry compared to plant sources, and some products utilize a combination of sources including both plants and microbes. The zero-calorie sweetener rebaudioside M (Reb M) –naturally occurring in stevia (*Stevia rebaudiana*) – is synthesized through bioconversion of Reb A to Reb M by bacteria, a multistep process requiring extensive purification (Okonkwo et al., 2024). Cell culture-based production has been established for a semisynthetic precursor of the anticancer drug Taxol® from Pacific yew (*Taxus brevifolia*), yet synthetic conversion is required for the last steps (Croteau et al., 2006). A close tobacco relative (*Nicotiana benthamiana*) offers an alternative expression platform, amenable to transient expression of heterologous genes including those involved in terpene biosynthesis (H. Kim et al., 2025). For example, *N. benthamiana* was engineered to express a range of psychoactive indolethylamines from plants, fungi, and animals as well as new-to-nature variants with therapeutic potential (Berman et al., 2026). However, low yields and modification of the heterologous terpene by native enzymes remain a barrier to tobacco-based production systems (Dong et al., 2013). Though efforts to generate tobacco plants with altered secondary metabolite profiles are ongoing (Bally et al., 2025), alternative plant expression systems are desirable. These challenges highlight the need to establish an economically viable, high-yielding, sustainable expression systems for complex specialized metabolites.

Tomato (*Solanum lycopersicum*) fruit has several desirable traits as a heterologous expression platform (Y. Li et al., 2018). First, triterpene synthases form sterol-based metabolites which function early in fruit formation (Chen et al., 2025), and abundant terpenes with the central carotenoid lycopene accumulate during ripening while other terpene classes are minimal (Davidovich-Rikanati et al., 2007). Second, distinct TPS paralogs function in the fruit while their counterparts execute critical functions in leaves (Ezquerro et al., 2023) permitting engineering of terpenes in the fruit without disrupting physiological roles in vegetative tissues. Third, the infrastructure is in place for propagation in both field as well as controlled environments enabling year-round production and processing of fruit into paste or juice that could be coopted for use in fruit-derived products. With a generation cycle time of 90 days, available robust genomic and genetic resources, and amenability to genetic engineering, it is feasible to design, build, and test a chassis tuned for production of terpenes in fruit. Tomato engineering for production has already been demonstrated for biofortification with the pro-vitamin A β-carotene (Fu et al., 2026), the dietary supplement γ-aminobutyric acid (GABA) (Nonaka et al., 2017; commercialized by Sanatech Seed Co., 2020), and anthocyanin antioxidants (Butelli et al., 2008; distributed by Norfolk Healthy Produce, a U.S. subsidiary of Norfolk Plant Sciences). An abundant native capacity suggests leverage for further engineering of specialized products in a dedicated chassis. For example, tomatoes produce MVA-derived steroidal glycoalkaloids (SGAs). While SGAs are natural antifeedants in the field (You & van Kan, 2021), they are not critical in controlled agricultural environments. Thus, directing metabolic flux away from this pathway late in fruit development has the potential to free up additional precursors that could be utilized for heterologous terpene production.

To design a terpene-depleted chassis in tomato fruits, we identified a comprehensive panel of genes for removal, including known terpene biosynthetic genes and candidate genes with the potential to interfere with heterologous terpene expression. Specifically, we profiled the expression of all genes encoding tomato TPSs, P450s, and UGTs in the ripening fruit. We generated combinatorial stable knockout lines targeting 23 genes that did not impact plant growth or reproduction. Metabolic profiling of the resulting chassis lines confirmed the roles of known terpene-related genes and extended our understanding through combinatorial knockouts, demonstrating that tomatoes tolerate the simultaneous loss of several terpene-related genes. Furthermore, these combinatorial knockouts showed modulation of native carbon reallocation toward endogenous sinks beneficial for a biosynthetic chassis.

## 2 Results and Discussion

### 2.1 Annotation of the M82 genome

With a distinct profile of specialized metabolites, and related genomic resources developed, M82–the cultivar used in this study–is an established reference and preferred background for engineering of metabolism and fruit traits (e.g. Fan et al., 2016). To identify candidate fruit-specific terpene-related genes, we generated six Oxford Nanopore Technologies (ONT) cDNA and 19 Illumina RNA-seq libraries from various *S. lycopersicum* M82 tissues and treatments (**Table S1**). These libraries were used to annotate the PacBio long read-generated M82 genome (Alonge et al., 2022). This annotation, SollycM82.v1, containing 32,804 high confidence protein-coding genes representing 62,359 gene models (**Table S2**) was used to guide targeted genome editing.

### 2.2 Identification of fruit-relevant terpene synthases

To engineer a fruit-specific terpene-depleted chassis, we first identified genes associated with terpene biosynthesis in M82 fruit. Previously curated tomato mono-, di-, and sesqui-terpene synthases in the Heinz1706 genome assembly (SL3.0) (Zhou & Pichersky, 2020b) were used to search the SollycM82.v1 protein and genome sequences. Differences in TPS gene repertoire between Heinz1706 and M82 primarily represent variations in copy number of mono-, di-, and sesqui-TPS genes present in biosynthetic clusters. These could represent differences in germplasm, genome assembly, or annotation (**Table S3**). A BLAST search using the previously characterized *TRITERPENE SYNTHASE1* (*TTS1*) and *TTS2* genes (Z. Wang et al., 2011) revealed three tandem *TTS* type gene models in M82. The SollycM82.v1 protein sequences of *TTS1* and *TTS2* are nearly identical (>97%) to previously published M82 *TTS1* and *TTS2* sequences (Z. Wang et al., 2011); the third *TTS* is ∼85% identical to *TTS1/TTS2* and was named *TTS3.* All three TTS sequences are also present in the Heinz1706 genome SL4.0. For tetra-TPSs, a BLAST search with *PHYTOENE SYNTHASE* (*PSY*) yielded all three previously reported *PSY* sequences (Ezquerro et al., 2023; Stauder et al., 2018); no additional *PSY* genes were identified. In summary, the complement of *PSY* and *TTS* genes in the M82 genome matches the Heinz1706 genome with limited differences in copy number of mono-, di-, and sesqui-TPS genes. This variation is consistent with previous reports that terpene biosynthesis genes are highly divergent, and many are dispensable across the tomato pan-genome (N. Li et al., 2023; X. Yu et al., 2022).

To determine which TPS genes may interfere with or compete with the production of heterologous terpenes in fruit, gene expression patterns were examined using a published fruit development gene expression atlas that included 11 tissues across 10 stages of ripeness from anthesis to red ripe (Shinozaki et al., 2018) and expression profiles from vegetative tissues and whole fruit generated in this study (**Figure 1; Table S4, Dataset 1**). *PSY1* is highly expressed in all fruit tissues beginning at the breaker stage through red ripe (**Figure 1**). Consistent with previous reports (Ezquerro et al., 2023; Stauder et al., 2018), *PSY2* and *PSY3* have low and no expression in fruit, respectively. *CrtR-b2* (encoding beta-carotene 3-hydroxylase), a downstream gene in the carotenoid pathway (Galpaz et al., 2006), showed a similar expression pattern to *PSY1* though at lower levels. The expression pattern of these carotenoid genes mirrors high expression of the well characterized *polygalacturonase* (PG) gene (Bird et al., 1988; Dellapenna et al., 1989; Montgomery et al., 1993; Nicholass et al., 1995) in total pericarp and septum beginning at the breaker stage. The three *TTS* genes have distinct expression patterns (**Figure 1**). *TTS1* is expressed at low levels in total pericarp, at higher levels in the outer epidermis but most strongly in leaf tissue. *TTS2* is expressed in the total pericarp, both the inner and outer epidermis with limited expression in the leaves, while *TTS3* is almost exclusively expressed in seeds. Of the putative mono- di-and sesqui-TPS genes, only SollycM82_v1.7G027980, a putative *ent*-kaurene synthase which is related to gibberellic acid metabolism (**Figure S1; Table S5**) has high expression in ripening fruit while five other mono-, di-, and sesqui-TPSs were expressed either very early in fruit development or in seeds.

**Figure 1.**
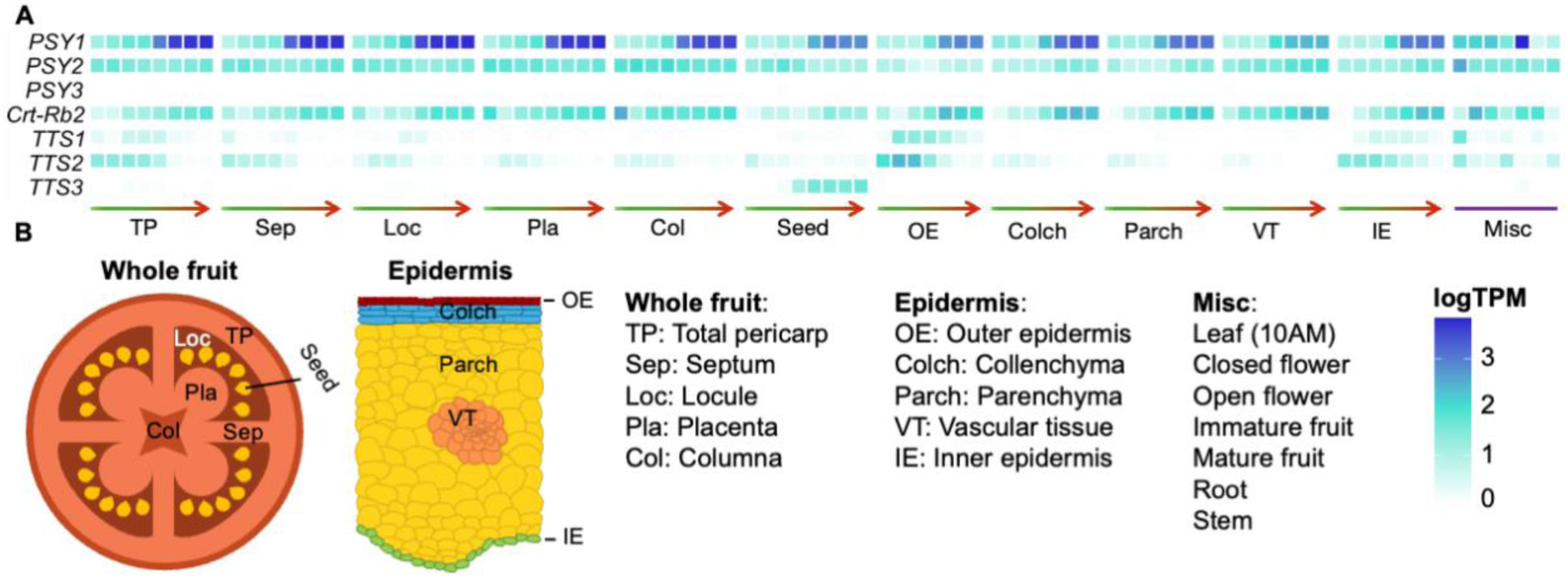
Expression of *PSY*, *TTS*, and *CrtR-b2* genes in fruit and select other developmental tissues. **(A)** Expression profiles of *PSY*, *TTS*, and *CrtR-b2* genes across an atlas of tomato fruit tissues **(B)** from anthesis to red ripe (green to red arrows) and other plant tissues (Misc).

For production of a heterologous terpene in tomato fruit, we hypothesize that the mono-, di-, and sesqui-TPS genes generally lack fruit expression and thus, would not impact fruit terpene production. However, *TTS2* is expressed in the total pericarp. Likewise, *PSY1* and *CrtR-b2* are strongly expressed in fruit and known to be responsible for producing abundant fruit carotenoids. Thus, we used gene editing technologies to create loss-of-function alleles of *TTS2*, *PSY1*, and *CrtR-b2* to not only eliminate production of native terpenes in the pericarp but also to free up a large precursor pool that can be diverted to production of desired heterologous terpenes.

### 2.3 Identifying fruit specific P450s and UGTs

In addition to TPSs that can divert metabolic resources away from desired products, P450s and UGTs have the potential to modify heterologous terpenes (Arendt et al., 2016). Putative P450 and UGT genes were identified in the M82 genome by Pfam designations PF00067 and PF00201, respectively, resulting in 349 putative P450s and 183 putative UGTs. P450s with an amino acid length less than 300 and UGTs less than 200 amino acids were excluded from further analyses resulting in 288 P450s with a median length of 503 amino acids and 176 UGTs with a median length of 470 amino acids. Tomato P450s were recently characterized in the Heinz1706 SL4.0 genome (Tang et al., 2024). Using Genespace (Lovell et al., 2022), we identified syntelogs of the Heinz1706 P450s in M82 (**Table S6**). For the 252 M82 P450s with a 1-to-1 syntelog relationship, we utilized the Heinz1706 names (**Table S6**); the 176 M82 UGTs were named based on location in the M82 genome (**Table S7**).

Both UGTs and P450s were filtered based on expression, requiring ≥50 transcripts per million (TPM) in at least one fruit expression dataset and a mean TPM ≥10 in the breaker through red ripe stages in the total pericarp (**Tables S8, S9**). To determine the evolutionary relationships of the 32 P450s and 17 UGTs that met these criteria, phylogenetic trees were constructed. Fruit-expressed P450s group into well-established clan groupings with 15 P450s in clan 71, the most abundant clan, as well as six P450s in clan 72, and 11 P450s in four other clans (**Figure 2A**). One of the fruit-expressed P450s, *CYP73A24a*, is duplicated in the M82 genome relative to Heinz1706, this duplicate was named *CYP73A24b*. Fruit-expressed UGTs fall into several clades with conservation within each group lower than the P450s (**Figure 2B**).

**Figure 2.**
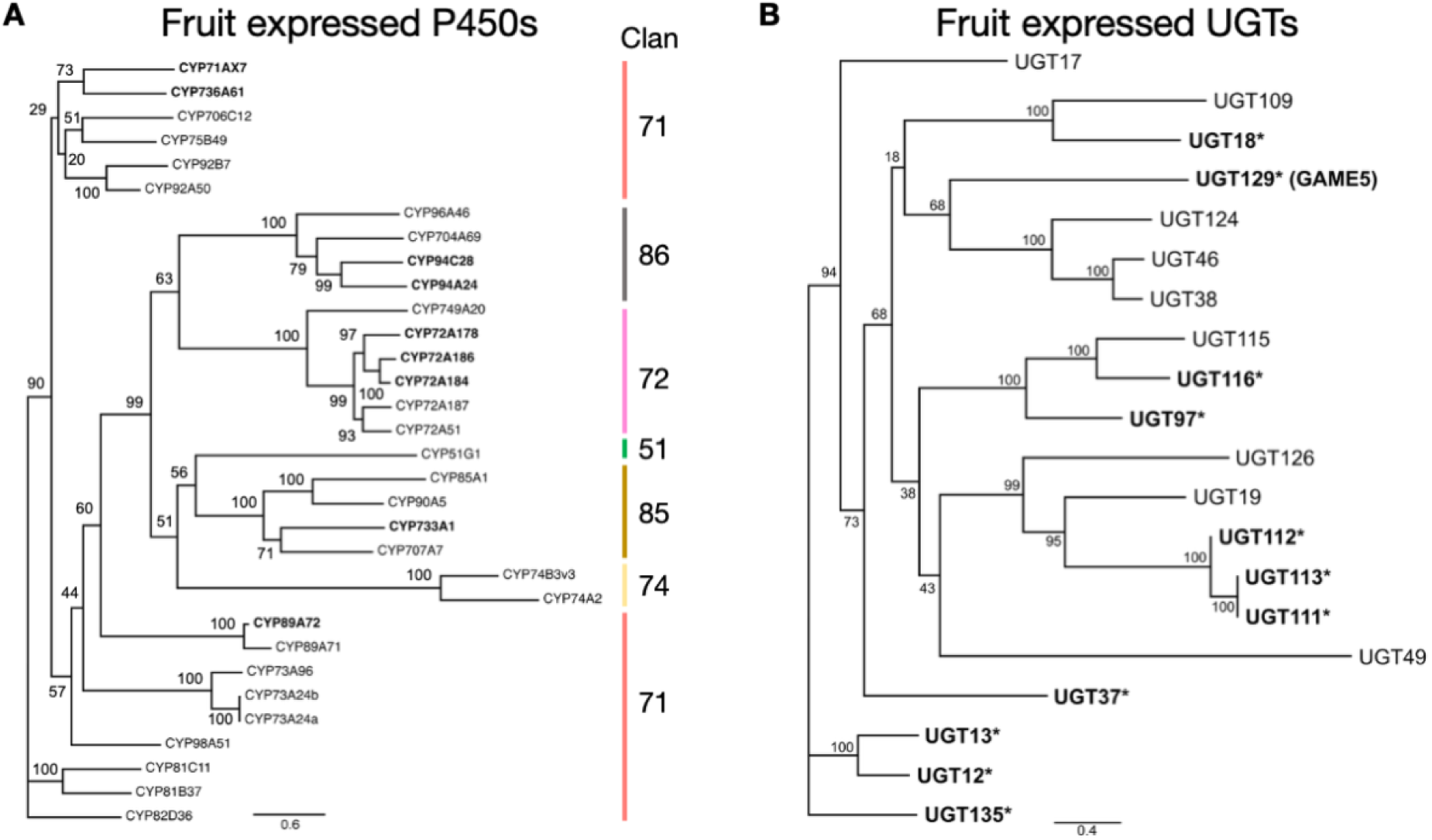
Fruit expressed P450 and UGT phylogenetic trees. Phylogenetic trees for fruit specific P450 with clan names **(A)** and UGT **(B)** proteins. Bold asterisks denote genes targeted for gene editing. Branch labels represent bootstrap values. Scale bars represent substitutions per site.

Using literature reports, phylogeny, and expression patterns, we further refined P450s and UGTs to generate a final list of candidates (genes with asterisks in **Figure 2**) yielding nine P450s and 11 UGTs that may confound expression of heterologous terpenes. Due to recent gene duplications, one UGT (SollycM82_v1.9G023450; *SlUGT112*) is nearly identical to two other UGTs (SollycM82_v1.9G023420; *SlUGT111* and SollycM82_v1.9G023470; *SlUGT113*) present in a local array thereby masking individual expression estimations. All three protein sequences are 100% identical across the first 223 amino acids but differ in 3’ exon structure; thus, all three UGTs were included in construction of our terpene-depleted chassis. One of the UGTs, *UGT129*, encodes GLYCOALKALOID METABOLISM5 (GAME5) which is involved in SGA biosynthesis (Szymański et al., 2020). The final list of genes targeted for gene editing includes two TPSs (*PSY1*, *TTS2)*, nine P450s, 11 UGTs, and *CrtR-b2* (**Table 1**, **Figure 3, Table S10**).

**Figure 3.**
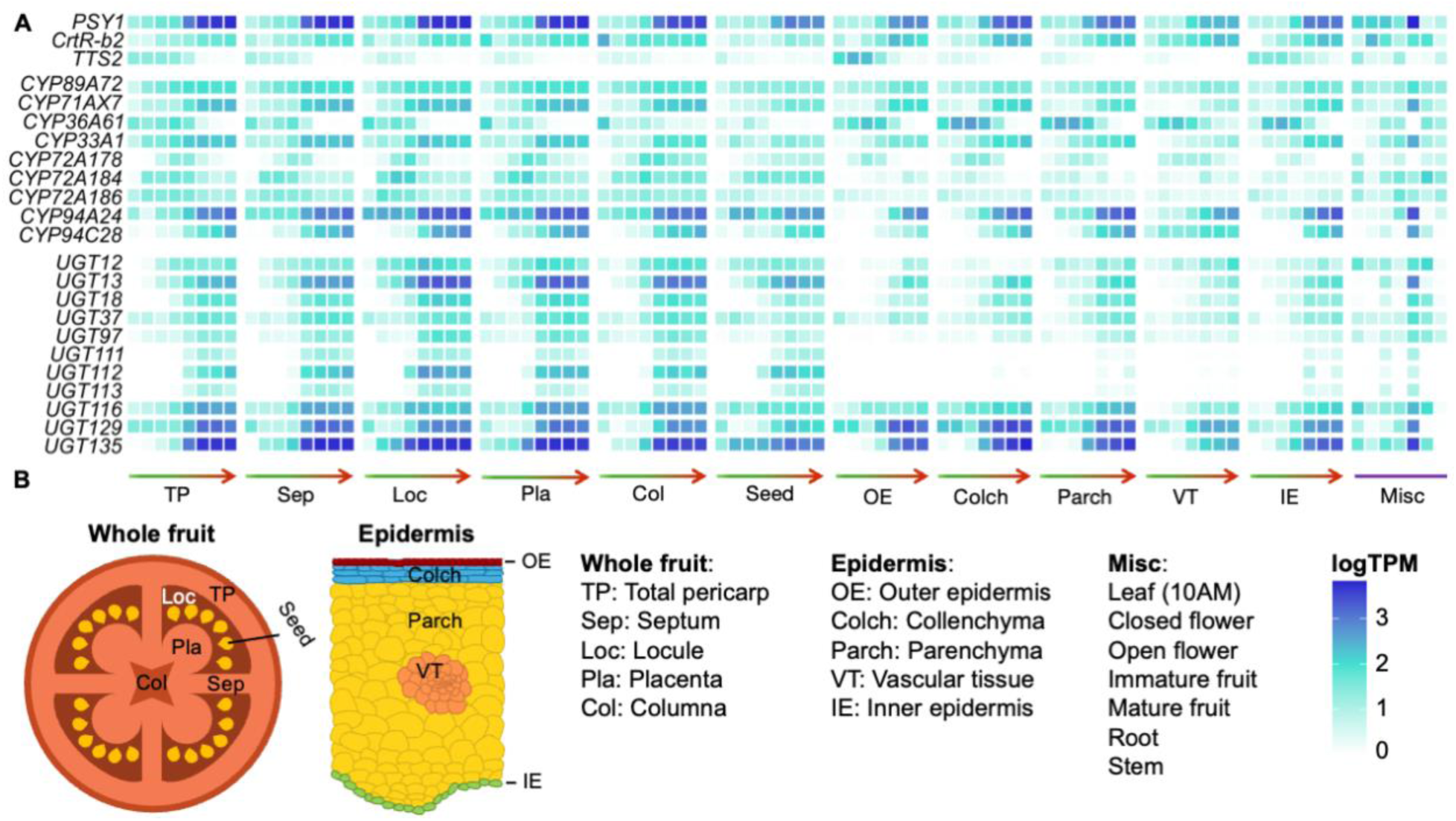
Expression of candidate terpene-related genes targeted for editing across fruit development and other organs. **(A)** Expression profiles of candidate genes across an atlas of tomato fruit tissues **(B)** from anthesis to red ripe (green to red arrows) and other plant tissues (Misc).

**Table 1.**
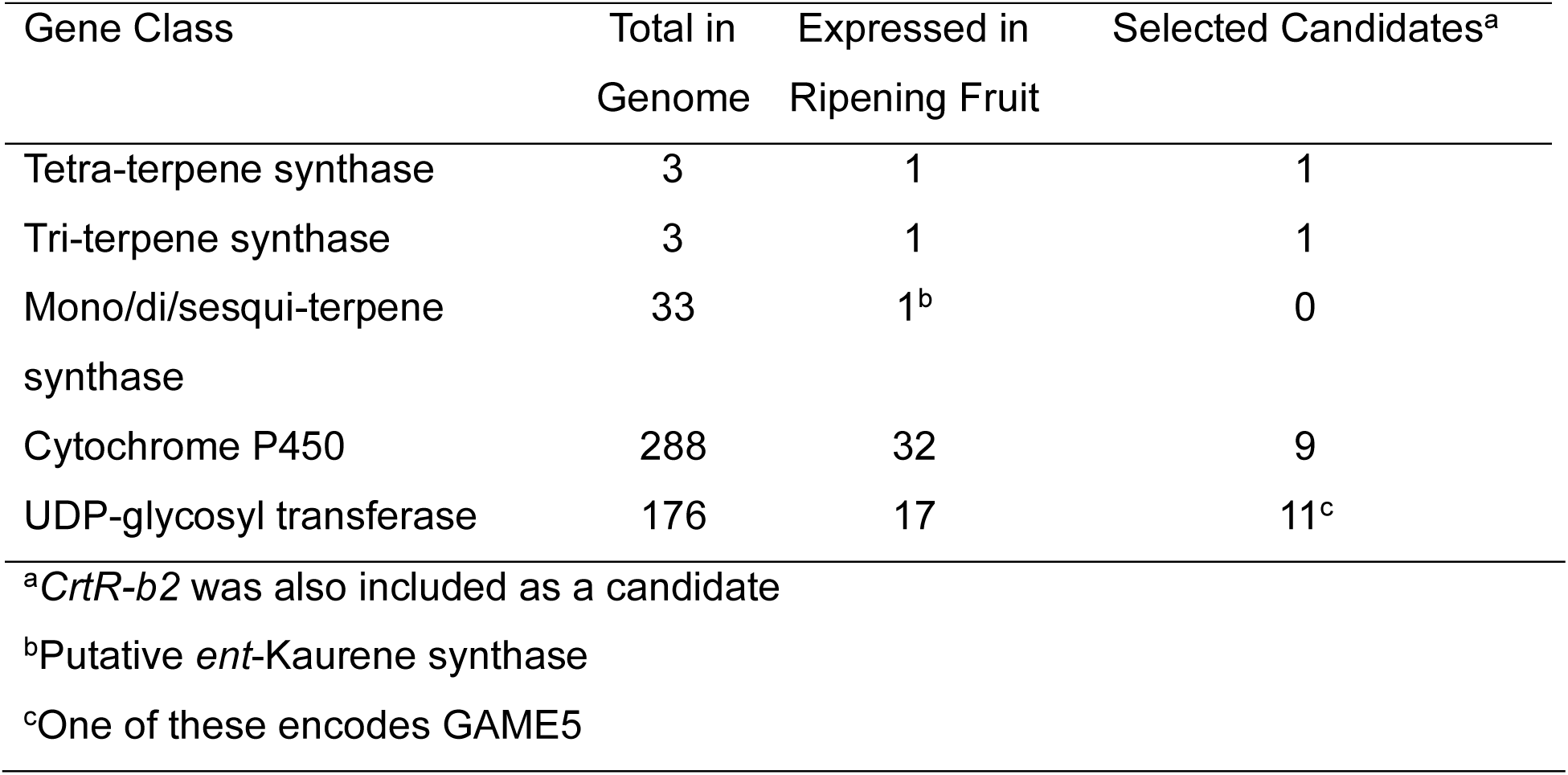
Genes evaluated and selected for construction of a terpene-depleted chassis in tomato fruit.

### 2.4 Combinatorial knockouts of candidate genes

To generate heritable knockouts of these terpene-related genes, we assembled four pairs of constructs targeting candidate genes based on chromosome location. For each set of target genes, two parallel constructs were made. One construct utilized the *35S* promoter to drive expression of *Cas9* and included *AmCyan* as a visual marker along with single guide RNAs (sgRNAs). The other construct used the *Ef1a* promoter and utilized luciferase as a reporter (**Table 2, Table S11**). pJPC716 and pJPC720 targeted three UGTs from chromosomes *1* and *2* and *TTS2* from chromosome 12; pJPC663 and pJPC722 targeted three P450s and a UGT from chromosomes *7* and *8;* pJPC717 and pJPC721 targeted *CrtR-b2*, *PSY1,* four P450s, and a UGT distributed across chromosomes *3* and *4*; and pJPC718 and pJPC723 targeted two P450s and six UGTs from chromosomes *9* and *10* (**Figure 4**). To maximize editing, two sgRNAs positioned near the beginning of the coding sequence were used for each gene. Due to sequence similarity, a single pair of sgRNAs was used for *SlUGT111*, *SlUGT112*, and *SlUGT113*. These constructs were transformed into M82, and a minimum of eight independent events were generated per line. We detected edits for all targets in at least one T_0_ plant from each construct (**Table 2, Figure S2**). Both promoters driving *Cas9* worked with similar efficiency.

**Figure 4.**
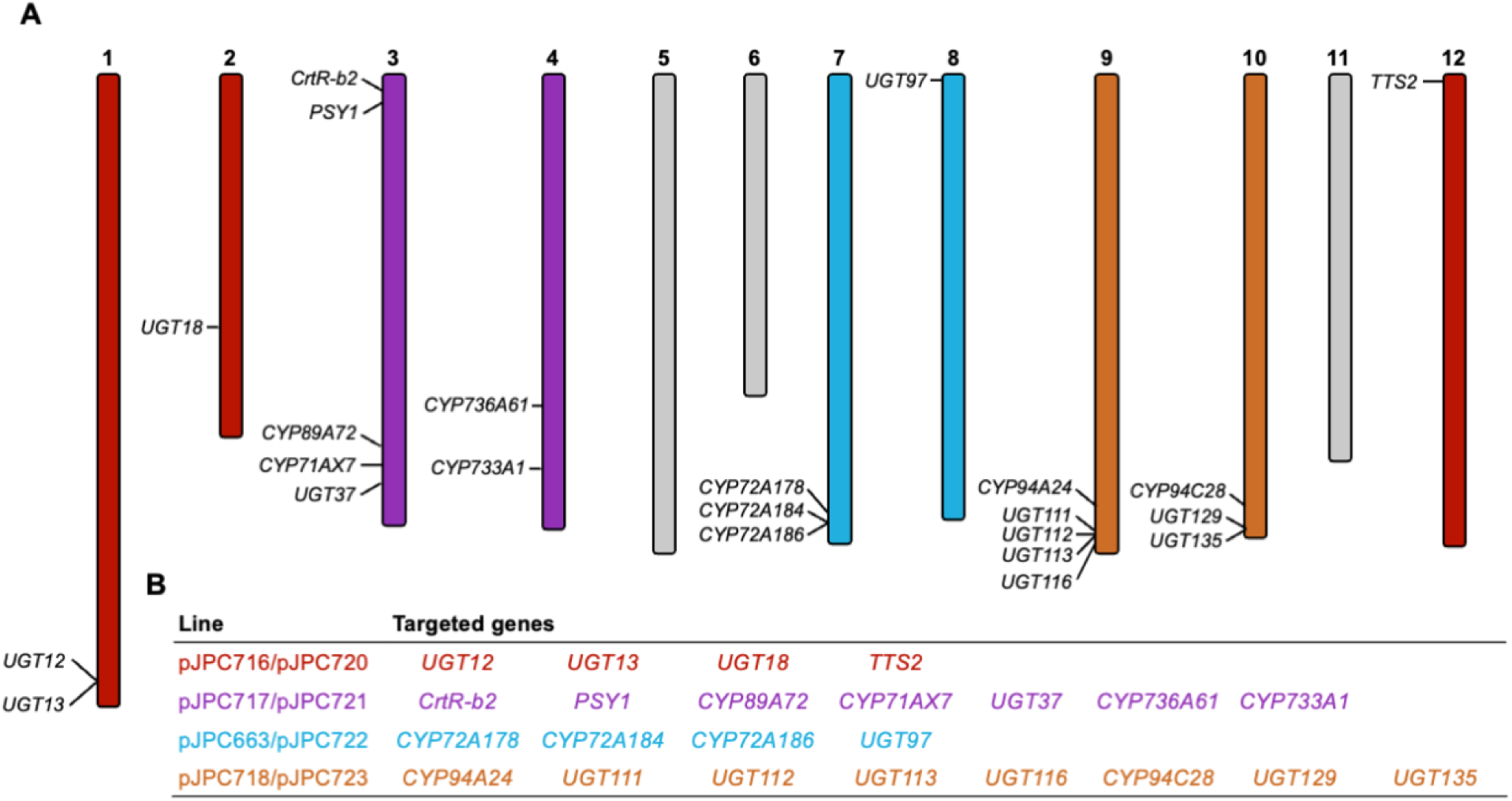
Genome distribution of candidates and construct organization. **(A)** Location of each candidate gene in the genome. **(B)** Constructs targeting groups of candidates by location.

**Table 2.** T_0_ editing outcomes.

| **Vector** | **CAS9 Promoter** | **Marker** | **# T<sub>0</sub> Plants** | **# CAS9+** | **# All Targets Edited** | **# All Biallelic** |
| --- | --- | --- | --- | --- | --- | --- |
| pJPC663 | 35S | AmCyan | 16 | 4 | 3 | 1 |
| pJPC716 | 35S | AmCyan | 15 | 5 | 1 | 0 |
| pJPC717 | 35S | AmCyan | 12 | 5 | 1 | 1 |
| pJPC718 | 35S | AmCyan | 26 | 11 | 1 | 0 |
| pJPC720 | Ef1a | Luc | 9 | 4 | 3 | 3 |
| pJPC721 | Ef1a | Luc | 8 | 4 | 1 | 1 |
| pJPC722 | Ef1a | Luc | 12 | 2 | 1 | 1 |
| pJPC723 | Ef1a | Luc | 24 | 15 | 1 | 1 |

Based on high expression of the target genes in ripe fruit, we prioritized three constructs (663, 721, and 723) and advanced individuals with maximal biallelic edits across 19 target loci to the T_3_ generation for downstream analyses. ONT long read genomic sequencing was performed on two individuals from each of the selected lines (663-5-3, 663-5-5, 721-16-1, 721-16-5, 723-4-1, and 723-4-2) to determine overall impacts of editing on genome sequence (**Table S12**). Of the 19 targets from these three lines, 18 targets had lesions predicted to disrupt function by producing a truncated protein in both T_3_ individuals sequenced (**Figure 5**). However, *SlCYP94A24* had an in-frame 36bp deletion. Loss of these 12 amino acids may impact the function of this protein, yet it is possible that it may retain some or all enzymatic activity. *SlUGT111*, *SlUGT112*, and *SlUGT113* each have 4bp deletions at the same sgRNA site.

**Figure 5:**
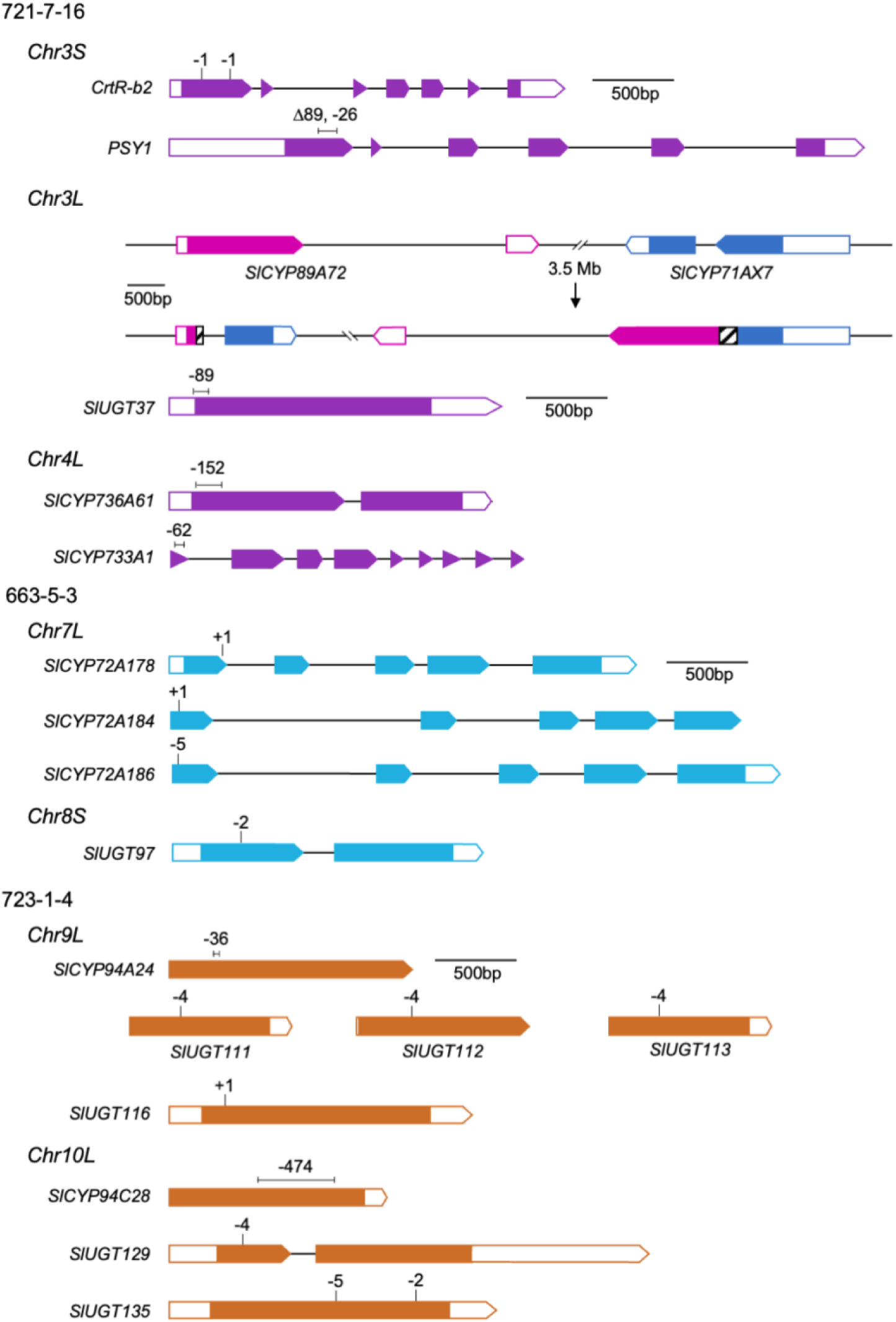
Lesions in terpene depleted chassis. Lesions in 19 candidate genes from three chassis. A 3.5Mb inversion between *SlCYP89A72* and *SlCYP71AX7* disrupts function of both genes (missing sequence hashed boxes results in frame shifts) leaving the intervening genes intact.

Because we simultaneously targeted several genomic locations, there was a possibility of chromosomal rearrangements. To assess genome structure, genome assemblies were generated from the long read ONT genomic DNA sequences and compared to the M82 genome (**Figure S3, Table S13**). The only structural variation observed was a 3.5Mb inversion between candidates SlCYP89A72 and SlCTP71AX7 on chromosome *3* in both sequenced pJPC721 T_3_ individuals. Maintaining intact genome structure will facilitate combining mutant lines and introducing heterologous expression pathways.

### 2.5 Metabolic changes were observed in combinatorial terpene depleted fruit

Untargeted analysis of terpene and terpene-derived pathways of the edited chassis showcased predicted changes in the red ripe fruit metabolome in a chassis-dependent manner (**Table S14**, **Figure 6**, **Datasets 2-7**). The carotenoid and steroidal glycoalkaloid pathways are two of the highest yielding terpene derived pathways in tomato fruit, and they were impacted significantly though differentially in the 663, 721, and 723 chassis (**Figure 6**; chemical structures in **Figure S4**). The 723 chassis carries a previously undescribed disruption of glycosylation in two steroidal glycoalkaloid pathway branches as shown by the depletion of escueloside A, dehydroescueloside A, and other derivatives. Depletion of the normally highly abundant products is accompanied by the accumulation of their respective precursors. These changes are consistent with a role for *UGT129* encoding GAME5 which has previously been indicated in the glycosylation of acetoxy-hydroxytomatine to form esculeoside A (Szymański et al., 2020). Considering the structural similarity between the two precursors, it is possible GAME5 can catalyze both glycosylations disrupted in the 723 chassis, forming escueloside A and dehydroescueloside A from acetoxy-hydroxytomatine and acetoxy-hydroxy-dehydrotomatine, respectively. These two relatively large sterol precursors differ only by the lack or presence of a carbon-carbon double bond (Sonawane et al., 2023; **Figure S4**).

**Figure 6.**
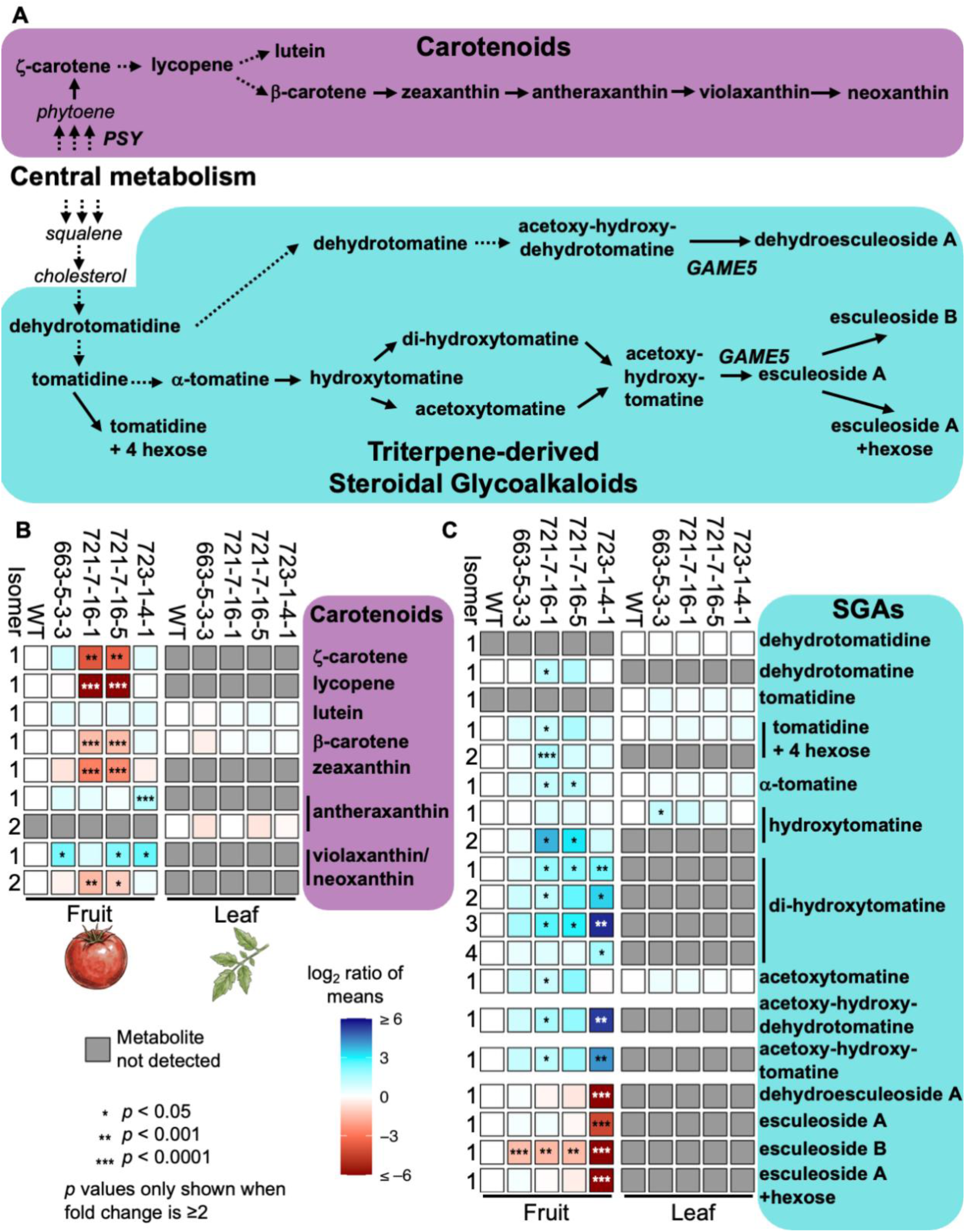
Metabolic changes in edited plants. **(A)** Key steps in carotenoid and steroidal glycoalkaloid (SGA) biosynthetic pathways. Bold metabolites were quantified in fruit and/or leaf while italicized compounds were not measured in either tissue. Relative abundance of **(B)** carotenoids and **(C)** SGAs in mutant lines compared to M82 (WT).

The metabolic impact of disrupted biosynthesis in the 721 chassis is dominated by the *psy1* mutation resulting in a broad loss of carotenoid-related metabolites in mutant individuals. However, we also observed an unexpected increase in abundance of various SGAs suggesting rerouting of metabolic resources like carbon, sugar, or isoprenoid precursors towards other metabolites. Hence, the 721 and 723 chassis metabolic profile supports expendable carbon for further genetic engineering. A subtle, but significant impact on the metabolite profile in the 663 mutant may provide leads to identify novel roles for yet uncharacterized genes targeted.

The observed distinct metabolite modulation in our lines provides support for known genes in the context of metabolically related knockouts. A possible interplay with other candidates may reveal previously unknown gene functions. Our approach also demonstrates that tomato can tolerate simultaneous loss of several terpenoid-related factors while still producing healthy fruit exhibiting robust physiology, a requirement for further metabolic engineering.

## 3 Conclusions

Alternative platforms for producing high-value plant chemicals will enable increased flexibility for biotechnology and economic viability for medicinal, nutritional, flavor, and other industrially relevant plant natural products. We present tomato as an agronomically viable chemical synthesis chassis for a variety of terpenoid natural products. Tomato has many desirable characteristics for a biosynthetic platform; however, heterologous products must compete with native pathways for metabolic resources. Combinatorial gene editing enabled us to simultaneously target known and putative genes to efficiently generate lines depleted for multiple enzymes related to specialized metabolism. We observed expected changes in metabolite abundances validating our approach to computationally predicting candidates which did not impact essential metabolic processes in the fruit indicating it is phenotypically robust with plasticity across families of the major terpene carbon sinks. Unexpected changes offer the potential to uncover novel biology masked by redundancy in large enzyme families. Taken together, our results support the development of tomato as an agronomically viable chemical synthesis chassis for a variety of terpenoid natural products.

## 4 Materials and Methods

### 4.1 Plant growth

For RNA isolation, M82 plants were grown in a growth chamber under a 25.5°C day/18°C night temperature regiment with 15-hour daylength of 300 µmol/m2/s light intensity. For fruit production, M82 were grown in a greenhouse at the University of Minnesota maintained within temperature ranges of 18.3°C-23.9°C day/17.8°C-23.9°C night with light levels > 920 µmol/m2/s or supplemented with 300 µmol/m2/s and 16-hour daylength.

### 4.2 M82 RNA isolation, library construction, and sequencing

A set of vegetative and reproductive tissues was collected in triplicate for transcriptome analyses (**Table S1**). At the flowering stage, a leaf time-course was collected sampling every four hours starting at 6am (dawn). For open and closed flowers, sepals were removed. Fruit were collected at immature and red ripe stages. Stem and root tissues were also collected from mature plants. For Methyl Jasmonate (MeJA) treatment, 250 µM MeJA was applied with a spray bottle until leaves were drenched and samples collected after 24 hours post-application. For heat treatment, plants were well watered to avoid drought stress prior to elevating the temperature to 37°C day and 28°C night; samples were collected after 24 hours. RNA was isolated using a modified hot borate protocol (Wan & Wilkins, 1994), treated with DNase using the Ambion Turbo DNA-free™ Kit (Thermo Fisher Scientific, Waltham, MA), quantitated using a Qubit and quality assessed using gel electrophoresis and NanoDrop. RNA-seq libraries were generated using the Illumina Stranded mRNA Prep Ligation kit with IDT for Illumina RNA UD Indexes and sequenced on an Illumina NovaSeq 6000 generating paired end 50 nt reads (Illumina, San Diego, CA). ONT cDNA libraries were generated from six RNA samples (**Table S1**) using the PCR-cDNA Sequencing Kit (SQK-PCS109) and sequenced on R9 FLO-MIN106 Rev D flow cells using a MinION Mk1B device.

### 4.3 M82 genome annotation

RNA-seq libraries (**Table S1**) were cleaned using Cutadapt (Martin, 2011) (v2.10) with a minimum length of 40 nt and quality cutoff of 10. Cleaned reads were aligned to the M82 genome using HISAT2 (D. Kim et al., 2019) (v2.1.0) with a maximum intron size of 5000 nt. Pychopper (v2.5.0; github.com/epi2me-labs/pychopper) was used to process the ONT cDNA reads; passed reads greater than 500 nt were aligned to the M82 genome using minimap2 (H. Li, 2018) (v2.17-r941) with a maximum intron length of 5,000 nt. Aligned RNA-seq and ONT cDNA reads were assembled using Stringtie (Kovaka et al., 2019) (v2.2.1); transcripts less than 500 nt were removed.

A repeat sequence library for M82 was constructed and used to mask the genome as described previously (Wood et al., 2026). Using the soft-masked M82 genome sequence with the aligned RNA-seq libraries as hints, BRAKER2 (Brůna et al., 2021; Hoff et al., 2019) (v2.1.6) was used to generate initial genome models which were refined using two rounds of PASA2 (Campbell et al., 2006; Haas et al., 2005) (v2.4.1) yielding the working gene model set. Gene models lacking expression evidence, or a PFAM domain match, or in which a partial gene model contained an interior stop codon were filtered out to yield a high confidence gene set. Functional annotation was assigned by searching the gene model proteins as described previously (Wood et al., 2026).

Candidate terpene-related genes were manually inspected for gene structure using transcript evidence from (1) mature and immature fruit bulk RNA-seq libraries that were aligned to the M82 genome using HISAT2 (options: -q --no-mixed --no-discordant --rna-strandness RF) (D. Kim et al., 2019) and (2) ONT cDNA alignments as described above. A subset of terpene-related gene models were manually curated to address misannotations.

### 4.4 Transcript quantification

A developmental time course of ripening M82 tomato fruit representing 483 RNA-seq 101 nt single-end datasets (Shinozaki et al., 2018), were downloaded from SRA PRJNA391024 (**Table S4**). A total of 36 50 nt paired-end datasets representing various whole tissues were generated in this study (available under SRA PRJNA753098). All datasets were quality trimmed and filtered using the bbduk function of BBTools (https://jgi.doe.gov/data-and-tools/software-tools/bbtools) using options: ftl=1 qtrim=lr trimq=20 maq=20 minlen=36. On average, 11.5 million and 36.8 million clean filtered reads remained in the PRJNA391024 and PRJNA753098 datasets, respectively.

Transcript abundance was determined for all datasets using kallisto (Bray et al., 2016) with options: PRJNA391024: --fr-stranded or --rf-stranded, --single -l 59 -s 1, PRJNA753098: --rf-stranded. Strand specificity of RNA-seq libraries was determined by comparing the kallisto mapping rate when reads were specified as either forward or reverse (--fr-stranded or --rf-stranded). All PRJNA753098 libraries are reverse stranded whereas PRJNA391024 includes a series of forward stranded libraries from laser microdissection samples while libraries generated from hand dissection samples were reverse stranded. Reads were pseudoaligned to high confidence representative gene models and quantified in transcripts per million (tpm).

### 4.5 Nomenclature of UGTs

UGTs were previously characterized using the SL2.5 version of the Heinz1706 genome (G. Yu et al., 2022) and were named by genomic location without regard to phylogeny. Thus, we also named M82 UGTs based on location in the M82 genome. As 128 M82 UGTs have a 1-to-1 syntelog with the previously named Heinz1706 UGTs, these names were also included in **Table S7**.

### 4.6 Phylogenetic trees

Full-length protein sequences from the SollycM82_v1 representative gene models were obtained for each fruit expressed P450 and UGT and aligned using MAFFT (v7.520 default options) (Katoh & Standley, 2013). Phylogenetic trees were built from each alignment by RAXML (v8.2.12; Stamatakis, 2014) with the PROTGAMMA AUTO model, algorithm a, and 1000 bootstraps. Trees were visualized using Geneious (v2023.2.1; Kearse et al., 2012).

### 4.7 Generating combinatorial knockout lines

The combinatorial knockout constructs were generated using a Golden Gate cloning system available from Addgene (https://www.addgene.org/browse/article/28244504/). Pairs of sgRNAs ∼100bp apart in the first or second exon of each target gene were cloned into level 0 vectors. See **Table S15** for sgRNA sequences and genotyping primers. Expression of each sgRNA was driven by an AtU6 promoter. Up to 4 pairs of sgRNAs were cloned into level 1 vectors and up to three sgRNA level 1 vectors were included in the final tDNA for a capacity of up to 24 sgRNAs. One set of constructs utilized the *35S* promoter to drive expression of CAS9 and included AmCyan as a visual marker while the other set used the *Ef1a* promoter and utilized luciferase as a reporter (**Table 2, Table S11**). Plasmid sequences can be found in **Supplemental files 1-8**. Sanger sequencing and Inference of CRISPR Edits (ICE CRISPR Analysis. 2025. v3.0. EditCo Bio) analysis was used to genotype each target in the resulting transgenic plants.

### 4.8 Metabolic analysis of fruit and leaves

#### 4.8.1 Plant tissue processing and metabolite extraction

Fruit and leaf tissue were frozen with liquid nitrogen, ground with a mortar and pestle, and lyophilized after harvesting. To extract carotenoids and other similarly fractionable metabolites, an organic solvent extraction from (Emiliani et al., 2018) using methanol, water, Tris-HCl solution, and chloroform was adopted with modifications. To extract steroidal glycoalkaloids and other similarly fractionable metabolites, an organic solvent extraction with acidified methanol and water was used.

For additional details on the plant tissue processing and metabolite extraction, please refer to Supplemental Methods.

#### 4.8.2 UPLC/MS analysis and ion characterization

All plant metabolite extracts were run on a Waters Xevo G2-XS Ultra Performance Liquid Chromatography (UPLC)/ Quadrupole Time of Flight (QToF) system (UPLC/QToF) in the Michigan State University Mass Spectroscopy and Metabolomics Core. UPLC/QToF data was filtered and analyzed using the peak picker software Progenesis QI (Nonlinear Dynamics, Version 3.0.7600.27622). Analyzed ions and genotype-specific patterns were screened for tomato metabolite isomers and their respective pathways. Isomer identifications with a mass error below 5 parts per million (ppm) were accepted. Accepted isomers and their relative, normalized abundances were subjected to two-tailed student’s t-tests assuming equal variance to provide statistical comparisons between genotypes.

For additional details on the UPLC/MS Analysis & Ion Characterization, please refer to Supplemental Methods.

## Author Contributions

C.R.B., B.R.H., D.F.V. planned and designed the study. N.C.D., J.C., L.R., J.P.H., C.S., L.P., J.C.W., and B.V. carried out the experimental work. N.C.D., J.C., and L.R. analyzed the data. N.C.D. wrote the first draft. All authors approve the manuscript.

## Acknowledgements

We acknowledge funding from the National Science Foundation (IOS- 2126592; C.R.B., B.R.H, and D.F.V.), Georgia Research Alliance (C.R.B.), Georgia Seed Development (C.R.B.), University of Georgia (C.R.B), and the National Institute of General Medical Sciences of the National Institutes of Health (T32-GM152798) (L.R.). We acknowledge the Texas A&M Genomics and Bioinformatics Service and the Michigan State Research Technology Support Facility for providing sequencing services. We acknowledge the University of Nebraska Plant Transformation facility for generating transgenic tomato. We acknowledge the Michigan State University Mass Spectrometry and Metabolomics Core for providing liquid chromatography and mass spectroscopy services We thank David Nelson (UTHSC) for validating and assisting with naming P450s. We thank Ellissa Rhuby (UMN) and Rishika Sahoo (UMN) for their assistance in genotyping.

## Data availability

The SollycM82.v1 annotation is available at spuddb.uga.edu/SollycM82_v1_download.shtml) as well as Figshare via doi (to be made public upon publication). SRA accession numbers for RNA-seq and WGS libraries generated or used in this study can be found in Tables S1 (genome annotation), S4 (expression profiling), and S12 (sequenced combinatorial mutants). The plasmids used in this study are available through Addgene (numbers in Table S11). Raw metabolomics data will be made available upon publication (Zenodo: doi 10.5281/zenodo.19713176).

## References

Akiel, M. A., Alshehri, O. Y., Aljihani, S. A., Almuaysib, A., Bader, A., Al-Asmari, A. I., Alamri, H. S., Alrfaei, B. M., & Halwani, M. A. (2022). Viridiflorol induces anti-neoplastic effects on breast, lung, and brain cancer cells through apoptosis. Saudi Journal of Biological Sciences, 29(2), 816–821. 10.1016/j.sjbs.2021.10.026

Alonge, M., Lebeigle, L., Kirsche, M., Jenike, K., Ou, S., Aganezov, S., Wang, X., Lippman, Z. B., Schatz, M. C., & Soyk, S. (2022). Automated assembly scaffolding using RagTag elevates a new tomato system for high-throughput genome editing. Genome Biology, 23(1), 258. 10.1186/s13059-022-02823-7

Arendt, P., Pollier, J., Callewaert, N., & Goossens, A. (2016). Synthetic biology for production of natural and new-to-nature terpenoids in photosynthetic organisms. In Plant Journal (Vol. 87, Number 1, pp. 16–37). Blackwell Publishing Ltd. 10.1111/tpj.13138

Aswathy, M., Vijayan, A., Daimary, U. D., Girisa, S., Radhakrishnan, K. V., & Kunnumakkara, A. B. (2022). Betulinic acid: A natural promising anticancer drug, current situation, and future perspectives. Journal of Biochemical and Molecular Toxicology, 36(12), e23206. 10.1002/jbt.23206

Bally, J., Jung, H., Mortimer, C., Naim, F., Philips, J. G., Hellens, R., Bombarely, A., Goodin, M. M., & Waterhouse, P. M. (2025). The Rise and Rise of Nicotiana benthamiana: A Plant for All Reasons. Annual Review of Phytopathology, 16, 42. 10.1146/annurev-phyto-080417

Berman, P., Höfer, J., Mehlman, H., Almekias-Siegl, E., Khersonsky, O., Dong, Y., Heinig, U., Sulimani, L., Kho Hao, L., Cohen, S., Peleg, Y., Meir, S., Rogachev, I., Meiri, D., Fleishman, S. J., & Aharoni, A. (2026). Complete biosynthesis of psychedelic tryptamines from three kingdoms in plants. Science Advances, 12, eaeb3034. https://www.science.org

Bird, C. R., Smith, C. J. S., Ray, J. A., Moureau, P., Bevan, M. W., Bird, A. S., Hughes, S., Morris, P. C., Grierson, D., & Schuch, W. (1988). The tomato polygalacturonase gene and ripening-specific expression in transgenic plants. Plant Molecular Biology, 11, 651–662.

Bray, N. L., Pimentel, H., Melsted, P., & Pachter, L. (2016). Near-optimal probabilistic RNA-seq quantification. Nature Biotechnology, 34(5), 525–527. 10.1038/nbt.3519

Brůna, T., Hoff, K. J., Lomsadze, A., Stanke, M., & Borodovsky, M. (2021). BRAKER2: Automatic eukaryotic genome annotation with GeneMark-EP+ and AUGUSTUS supported by a protein database. NAR Genomics and Bioinformatics, 3(1), 1. 10.1093/nargab/lqaa108

Butelli, E., Titta, L., Giorgio, M., Mock, H. P., Matros, A., Peterek, S., Schijlen, E. G. W. M., Hall, R. D., Bovy, A. G., Luo, J., & Martin, C. (2008). Enrichment of tomato fruit with health-promoting anthocyanins by expression of select transcription factors. Nature Biotechnology, 26(11), 1301–1308. 10.1038/nbt.1506

Campbell, M. A., Haas, B. J., Hamilton, J. P., Mount, S. M., & Robin, C. R. (2006). Comprehensive analysis of alternative splicing in rice and comparative analyses with Arabidopsis. BMC Genomics, 7. 10.1186/1471-2164-7-327

Chen, J., Liu, M., Zhang, Y., & Bai, F. (2025). Dissecting the biosynthesis, regulation, and metabolic engineering of steroidal glycoalkaloids in tomato. Journal of Integrative Plant Biology, jipb.70077. 10.1111/jipb.70077

Croteau, R., Ketchum, R. E. B., Long, R. M., Kaspera, R., & Wildung, M. R. (2006). Taxol biosynthesis and molecular genetics. In Phytochemistry Reviews (Vol. 5, Number 1, pp. 75–97). 10.1007/s11101-005-3748-2

Davidovich-Rikanati, R., Sitrit, Y., Tadmor, Y., Iijima, Y., Bilenko, N., Bar, E., Carmona, B., Fallik, E., Dudai, N., Simon, J. E., Pichersky, E., & Lewinsohn, E. (2007). Enrichment of tomato flavor by diversion of the early plastidial terpenoid pathway. Nature Biotechnology, 25(8), 899–901. 10.1038/nbt1312

de Matos Balsalobre, N., dos Santos, E., Mariano dos Santos, S., Arena, A. C., Konkiewitz, E. C., Ziff, E. B., Nazari Formagio, A. S., & Leite Kassuya, C. A. (2023). Potential anti-arthritic and analgesic properties of essential oil and viridiflorol obtained from *Allophylus edulis* leaves in mice. Journal of Ethnopharmacology, 301, 115785. 10.1016/j.jep.2022.115785

Dellapenna, D., Lincoln, J. E., Fischer, R. L., & Bennett, A. B. (1989). Transcriptional analysis of polygalacturonase and other ripening associated genes in Rutgers, *rin*, *nor*, and *Nr* tomato fruit. Plant Physiol, 90, 1372–1377. https://academic.oup.com/plphys/article/90/4/1372/6082280

DNP. (n.d.). Dictionary of Natural Products. In https://dnp.chemnetbase.com*: 34.2* (Number Accessed March, 2026). CRC Press, Taylor & Francis Group.

Dong, L., Miettinen, K., Goedbloed, M., Verstappen, F. W. A., Voster, A., Jongsma, M. A., Memelink, J., Krol, S. van der, & Bouwmeester, H. J. (2013). Characterization of two geraniol synthases from Valeriana officinalis and Lippia dulcis: Similar activity but difference in subcellular localization. Metabolic Engineering, 20, 198–211. 10.1016/j.ymben.2013.09.002

Emiliani, J., D’Andrea, L., Falcone Ferreyra, M. L., Maulión, E., Rodriguez, E., Rodriguez-Concepción, M., & Casati, P. (2018). A role for β,β-xanthophylls in Arabidopsis UV-B photoprotection. Journal of Experimental Botany, 69(20), 4921–4933. 10.1093/jxb/ery242

Ezquerro, M., Burbano-Erazo, E., & Rodriguez-Concepcion, M. (2023). Overlapping and specialized roles of tomato phytoene synthases in carotenoid and abscisic acid production. Plant Physiology, 193(3), 2021–2036. 10.1093/plphys/kiad425

Fan, P., Miller, A. M., Schilmiller, A. L., Liu, X., Ofner, I., Jones, A. D., Zamir, D., & Last, R. L. (2016). In vitro reconstruction and analysis of evolutionary variation of the tomato acylsucrose metabolic network. Proceedings of the National Academy of Sciences of the United States of America, 113(2), E239–E248. 10.1073/pnas.1517930113

Fu, J., Tieman, D., & Rathinasabapathi, B. (2026). Expressing CaCCS encoding capsanthin/capsorubin synthase in tomato increases fruit provitamin A content and yield. Plant Physiology, 200(3), 1–18. 10.1093/plphys/kiag062

Galpaz, N., Ronen, G., Khalfa, Z., Zamir, D., & Hirschberg, J. (2006). A chromoplast-specific carotenoid biosynthesis pathway is revealed by cloning of the tomato white-flower locus. Plant Cell, 18(8), 1947–1960. 10.1105/tpc.105.039966

Haas, B. J., Wortman, J. R., Ronning, C. M., Hannick, L. I., Smith, R. K., Maiti, R., Chan, A. P., Yu, C., Farzad, M., Wu, D., White, O., & Town, C. D. (2005). Complete reannotation of the *Arabidopsis* genome: Methods, tools, protocols and the final release. BMC Biology, 3, 7. 10.1186/1741-7007-3-7

Hoff, K. J., Lomsadze, A., Borodovsky, M., & Stanke, M. (2019). Whole-genome annotation with BRAKER. In Methods in Molecular Biology (Vol. 1962, pp. 65–95). Humana Press Inc. 10.1007/978-1-4939-9173-0_5

Katoh, K., & Standley, D. M. (2013). MAFFT Multiple Sequence Alignment Software version 7: improvements in performance and usability. Molecular Biology and Evolution, 30(4), 772–780. 10.1093/molbev/mst010

Kearse, M., Moir, R., Wilson, A., Stones-Havas, S., Cheung, M., Sturrock, S., Buxton, S., Cooper, A., Markowitz, S., Duran, C., Thierer, T., Ashton, B., Meintjes, P., & Drummond, A. (2012). Geneious Basic: an integrated and extendable desktop software platform for the organization and analysis of sequence data. Bioinformatics, 28(12), 1647–1649. 10.1093/bioinformatics/bts199

Kim, D., Paggi, J. M., Park, C., Bennett, C., & Salzberg, S. L. (2019). Graph-based genome alignment and genotyping with HISAT2 and HISAT-genotype. Nature Biotechnology, 37, 907–915. 10.1038/s41587-019-0201-4

Kim, H., Liu, L., Han, L., Park, K., Kim, H. J., Nguyen, T., Nazarenus, T. J., Cahoon, R. E., Haslam, R. P., Ciftci, O., Napier, J. A., & Cahoon, E. B. (2025). Oilseed-based metabolic engineering of astaxanthin and related ketocarotenoids using a plant-derived pathway: Lab-to-field-to-application. Plant Biotechnology Journal, 23(8), 3451–3464. 10.1111/pbi.70148

Kovaka, S., Zimin, A. V, Pertea, G. M., Razaghi, R., Salzberg, S. L., & Pertea, M. (2019). Transcriptome assembly from long-read RNA-seq alignments with StringTie2. Genome Biology, 20, 278. 10.1186/s13059-019-1910-1

Lanier, E. R., Andersen, T. B., & Hamberger, B. (2023). Plant terpene specialized metabolism: complex networks or simple linear pathways? Plant Journal, 114(5), 1178–1201. 10.1111/tpj.16177

Li, H. (2018). Minimap2: Pairwise alignment for nucleotide sequences. Bioinformatics, 34(18), 3094–3100. 10.1093/bioinformatics/bty191

Li, N., He, Q., Wang, J., Wang, B., Zhao, J., Huang, S., Yang, T., Tang, Y., Yang, S., Aisimutuola, P., Xu, R., Hu, J., Jia, C., Ma, K., Li, Z., Jiang, F., Gao, J., Lan, H., Zhou, Y., … Yu, Q. (2023). Super-pangenome analyses highlight genomic diversity and structural variation across wild and cultivated tomato species. Nature Genetics, 55(5), 852–860. 10.1038/s41588-023-01340-y

Li, Y., Wang, H., Zhang, Y., & Martin, C. (2018). Can the world’s favorite fruit, tomato, provide an effective biosynthetic chassis for high-value? metabolites? In Plant Cell Reports (Vol. 37, Number 10, pp. 1443–1450). Springer Verlag. 10.1007/s00299-018-2283-8

Lou, H., Li, H., Zhang, S., Lu, H., & Chen, Q. (2021). A review on preparation of betulinic acid and its biological activities. Molecules, 26(18), 5583. 10.3390/molecules26185583

Lovell, J. T., Sreedasyam, A., Schranz, M. E., Wilson, M., Carlson, J. W., Harkess, A., Emms, D., Goodstein, D. M., & Schmutz, J. (2022). GENESPACE tracks regions of interest and gene copy number variation across multiple genomes. ELIFE, 11,:e78526.

Mahroug, S., Burlat, V., & St-Pierre, B. (2007). Cellular and sub-cellular organisation of the monoterpenoid indole alkaloid pathway in *Catharanthus roseus*. Phytochemistry Reviews, 6(2–3), 363–381. 10.1007/s11101-006-9017-1

Martin, M. (2011). Cutadapt removes adapter sequences from high-throughput sequencing reads. EMBnet.Journal, 17(1), 10–12.

Montgomery, J., Pollard, V., Deikman, J., & Fischer, R. L. (1993). Positive and negative regulatory regions control the spatial distribution of polygalacturonase transcription in tomato fruit. The Plant Cell, 5(9), 1049–1062. https://about.jstor.org/terms

Mu, H., Sun, Y., Yuan, B., & Wang, Y. (2023). Betulinic acid in the treatment of breast cancer: Application and mechanism progress. Fitoterapia, 169, 105617. 10.1016/j.fitote.2023.105617

Nicholass, F. J., Smith, C. J. S., Schuch, W., Bird, C. R., & Grierson, D. (1995). High levels of ripening-specific reporter gene expression directed by tomato fruit polygalacturonase gene-flanking regions. Plant Molecular Biology, 28, 423–435.

Nonaka, S., Arai, C., Takayama, M., Matsukura, C., & Ezura, H. (2017). Efficient increase of γ-aminobutyric acid (GABA) content in tomato fruits by targeted mutagenesis. Scientific Reports, 7, 7057. 10.1038/s41598-017-06400-y

Okonkwo, C. E., Adeyanju, A. A., Onyeaka, H., Nwonuma, C. O., Olaniran, A. F., Alejolowo, O. O., Inyinbor, A. A., Oluyori, A. P., & Zhou, C. (2024). A review on rebaudioside M: The next generation steviol glycoside and noncaloric sweetener. In Journal of Food Science (Vol. 89, Number 11, pp. 6946–6965). John Wiley and Sons Inc. 10.1111/1750-3841.17401

Shinozaki, Y., Nicolas, P., Fernandez-Pozo, N., Ma, Q., Evanich, D. J., Shi, Y., Xu, Y., Zheng, Y., Snyder, S. I., Martin, L. B. B., Ruiz-May, E., Thannhauser, T. W., Chen, K., Domozych, D. S., Catalá, C., Fei, Z., Mueller, L. A., Giovannoni, J. J., & Rose, J. K. C. (2018). High-resolution spatiotemporal transcriptome mapping of tomato fruit development and ripening. Nature Communications, 9(1), 364. 10.1038/s41467-017-02782-9

Shukal, S., Chen, X., & Zhang, C. (2019). Systematic engineering for high-yield production of viridiflorol and amorphadiene in auxotrophic *Escherichia coli*. Metabolic Engineering, 55, 170–178. 10.1016/j.ymben.2019.07.007

Sonawane, P. D., Gharat, S. A., Jozwiak, A., Barbole, R., Heinicke, S., Almekias-Siegl, E., Meir, S., Rogachev, I., Connor, S. E. O., Giri, A. P., & Aharoni, A. (2023). A BAHD-type acyltransferase concludes the biosynthetic pathway of non-bitter glycoalkaloids in ripe tomato fruit. Nature Communications, 14(1), 4540. 10.1038/s41467-023-40092-5

Stamatakis, A. (2014). RAxML version 8: a tool for phylogenetic analysis and post-analysis of large phylogenies. Bioinformatics, 30(9), 1312–1313. 10.1093/bioinformatics/btu033

Stauder, R., Welsch, R., Camagna, M., Kohlen, W., Balcke, G. U., Tissier, A., & Walter, M. H. (2018). Strigolactone levels in dicot roots are determined by an ancestral symbiosis-regulated clade of the PHYTOENE SYNTHASE gene family. Frontiers in Plant Science, 9. 10.3389/fpls.2018.00255

Szymański, J., Bocobza, S., Panda, S., Sonawane, P., Cárdenas, P. D., Lashbrooke, J., Kamble, A., Shahaf, N., Meir, S., Bovy, A., Beekwilder, J., Tikunov, Y., Romero de la Fuente, I., Zamir, D., Rogachev, I., & Aharoni, A. (2020). Analysis of wild tomato introgression lines elucidates the genetic basis of transcriptome and metabolome variation underlying fruit traits and pathogen response. Nature Genetics, 52(10), 1111–1121. 10.1038/s41588-020-0690-6

Tang, M., Zhang, W., Lin, R., Li, L., He, L., Yu, J., & Zhou, Y. (2024). Genome-wide characterization of cytochrome P450 genes reveals the potential roles in fruit ripening and response to cold stress in tomato. Physiologia Plantarum, 176(3). 10.1111/ppl.14332

Tholl, D. (2015). Biosynthesis and Biological Functions of Terpenoids in Plants. In J. Schrader & J. Bohlmann (Eds.), Biotechnology of Isoprenoids. Advances in Biochemical Engineering/Biotechnology (Vol. 148, pp. 63–106). Springer. 10.1007/10_2014_295

Wan, C.-Y., & Wilkins, T. A. (1994). A modified hot borate method significantly enhances yield of high-quality RNA from cotton (*Gossypium hirsutum* L.). Analytical Biochemistry, 223, 7–12.

Wang, J., Li, S., Xiong, Z., & Wang, Y. (2016). Pathway mining-based integration of critical enzyme parts for de novo biosynthesis of steviolglycosides sweetener in Escherichia coli. In Cell Research (Vol. 26, Number 2, pp. 258–261). Nature Publishing Group. 10.1038/cr.2015.111

Wang, Z., Guhling, O., Yao, R., Li, F., Yeats, T. H., Rose, J. K. C., & Jetter, R. (2011). Two oxidosqualene cyclases responsible for biosynthesis of tomato fruit cuticular triterpenoids. Plant Physiology, 155(1), 540–552. 10.1104/pp.110.162883

Wood, J. C., Hamilton, J. P., Vaillancourt, B., Brose, J., Edger, P. P., & Buell, C. R. (2026). Chromosome-scale genome assembly for yellow wood sorrel, Oxalis stricta. G3: Genes, Genomes, Genetics, 16(3), jkaf317. 10.1093/g3journal/jkaf317

You, Y., & van Kan, J. A. L. (2021). Bitter and sweet make tomato hard to (b)eat. New Phytologist, 230(1), 90–100. 10.1111/nph.17104

Yu, G., Chen, Q., Chen, F., Liu, H., Lin, J., Chen, R., Ren, C., Wei, J., Zhang, Y., Yang, F., & Sheng, Y. (2022). Glutathione promotes degradation and metabolism of residual fungicides by inducing UDP-glycosyltransferase genes in tomato. Frontiers in Plant Science, 13, 893508. 10.3389/fpls.2022.893508

Yu, X., Qu, M., Shi, Y., Hao, C., Guo, S., Fei, Z., & Gao, L. (2022). Chromosome-scale genome assemblies of wild tomato relatives *Solanum habrochaites* and *Solanum galapagense* reveal structural variants associated with stress tolerance and terpene biosynthesis. Horticulture Research, 9, uhac139. 10.1093/hr/uhac139

Zhang, Y., Long, Y., Yu, S., Li, D., Yang, M., Guan, Y., Zhang, D., Wan, J., Liu, S., Shi, A., Li, N., & Peng, W. (2021). Natural volatile oils derived from herbal medicines: A promising therapy way for treating depressive disorder. Pharmacological Research, 164, 105376. 10.1016/j.phrs.2020.105376

Zhou, F., & Pichersky, E. (2020a). More is better: the diversity of terpene metabolism in plants. Current Opinion in Plant Biology, 55, 1–10. 10.1016/j.pbi.2020.01.005

Zhou, F., & Pichersky, E. (2020b). The complete functional characterisation of the terpene synthase family in tomato. New Phytologist, 226(5), 1341–1360. 10.1111/nph.16431

