## Supplementary Figures for "Engineering carotenoid and steroidal glycoalkaloid depleted tomato fruit for heterologous production of high value terpenes"

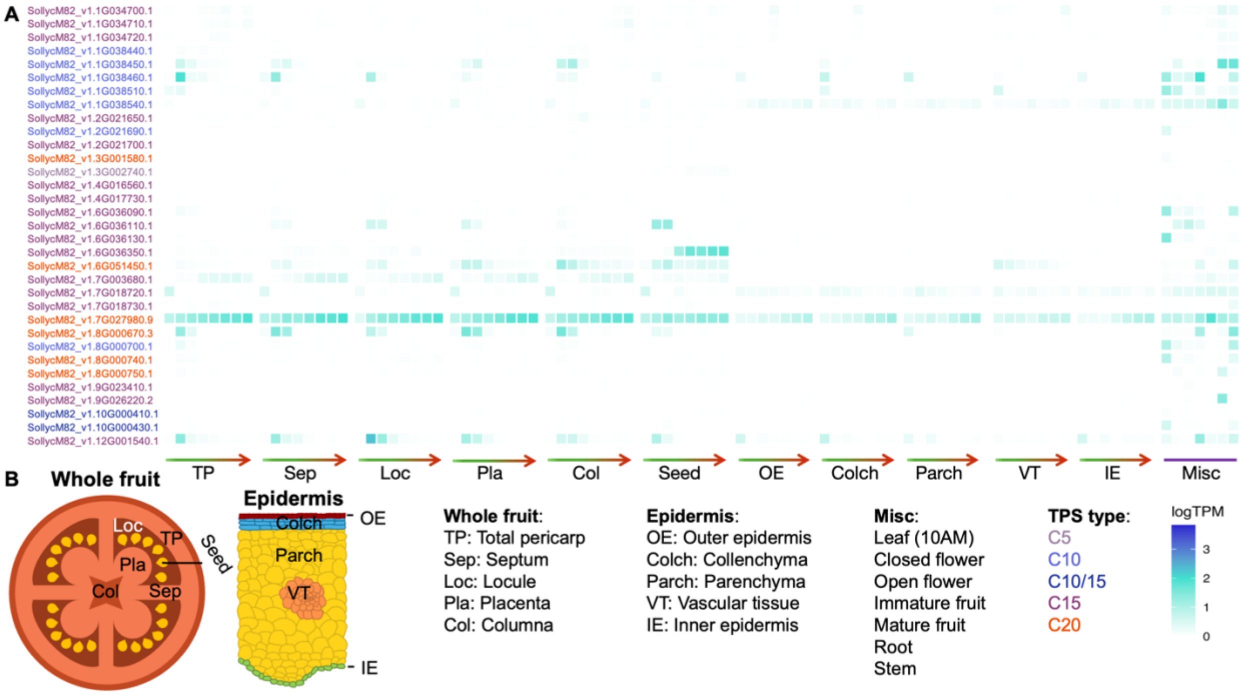


**Figure S1**. Mono-, di-, and sesquiterpene synthase expression in fruit.

**(A)** Expression profiles of mono, di and sesqui-terpene synthases across an atlas of tomato fruit tissues **(B)** from anthesis to red ripe (green to red arrows) and other plant tissues (Misc).

### Lesions in targeted genes

#### pJPC721

```
PSY1      MSVALLWVVSPCDVSNGTSFMESVREGNRFFDSSRHRNLVSNERINRGGGKQTNNGRKFS
mut       MSVALLWVVSPCDVSNGTSFMESVREGNRFFDSSRHRNLVSNERINRGGGKQTNNGRKFS

          VRSAILATPSGERTMTSEQMVYDVVLRQAALVKRQLRSTNELEVKPDIPIPGNLGLLSEA
          VRSAILATPSGVQCVYRLIQMYLDISCNK*

          YDRCGEVCAEYAKTFNLGTMLMTPERRRAIWAIYVWCRRTDELVDGPNASYITPAALDRW

          ENRLEDVFNGRPFDMLDGALSDTVSNFPVDIQPFRDMIEGMRMDLRKSRYKNFDELYLYC

          YYVAGTVGLMSVPIMGIAPESKATTESVYNAALALGIANQLTNILRDVGEDARRGRVYLP

          QDELAQAGLSDEDIFAGRVTDKWRIFMKKQIHRARKFFDEAEKGVTELSSASRFPVWASL

          VLYRKILDEIEANDYNNFTKRAYVSKSKKLIALPIAYAKSLVPPTKTASLQR*
```

```
CrtR-b2   MAAGISASASSRTIRLRHNPFLSPKSASTAPPVLFFSPLTRNFGAILLSRRKPRLAVCFV
mut       MAAGISASASSRTIRLRHNPFLSPKSA*

          LENEKLNSTIESESEVIEDRIQVEINEEKSLAASWLAEKLARKKSERFTYLVAAVMSSLG

          ITSMAILAVYYRFSWQMEGGEVPFSEMLATFTLSFGAAVGMEYWARWAHRALWHASLWHM

          HESHHRPREGPFEMNDVFAITNAVPAIALLSYGFFHKGIVPGLCFGAGLGITVFGMAYMF

          VHDGLVHKRFPVGPIANVPYFRRVAAAHQLHHSDKFDGVPYGLFLGPKELEEVGGLEELE

          KEVNRRIKISKGLL*
```

```
SlCYP89A72  MEIWFIVVVTLCVAFWLKSICNIIVSPNSNTKRNLPPGPYSFPVIGSLLWAKRTFSDLEP
mut         MEIWFIVVVTLCVAFWLKSICNIIVSPLQRDSLKAILLVI*

            ILRDLKAKYGPLITLNIGSRSAIFVASHSLAYQALVQQGAIFSNRPKAGPTSAVVNSNQR

            NISSAPYGPIWRLLRRNLTSEILHPSRIKSYSKARSWVLGILIQQLRNAQVDSVKLIDHF

            QYAMFCLLVLMCFGDKLEETQIRQIENIQRKLLLGFTRFNIINIFPRVGKIIFRNRWKEL

            IELRQEQESIIIPLIEARSRAREQRTEHGDEFVVAYVDTLLNLEFPEEKRNLNHGEIVTL

            CSEFLSAGTDTTSTALQWIMANLVKNPSIQEKLYQEIATVVGEKQSKLTDEEVVKEEDLQ

            KMPYLKAVILEGLRRHPPGHFVLPHTVTEEVELNGYVIPKDVTINFMVAEMGLDPKVWED

            PLEFRPERFLVEGSDNEGFDITGSREIKMMPFGAGRRICPAYALAMLHLEFFVANLIWHF

            QWKPVEGDDVDLTEILEFTVVMKNPLRARICPRVNSV*
```

```
SlCYP71AX7  MEFPWYSVVVPLIVFLFFLHNCLFTIFNTSNKKLPPSPRKLPIIGNLYQLGLHPHRSLYK
mut         MEFPWYSVVVPLIVFLFFLHNCLFTIFNTSNKKLPPSPRKLPIIGNLYQLGLHPHRSLYK

            LSKKYGPVMLLHLGSKPVLVASSVDAARDILKTHDLVWSTRPKSSIADGLLYGSKGVAFS
            LSKKYGPVMLLHLGSKPVLLPLLMLLAIS*

            NYSEYWRQVRSVIVLHLLSNKRVQSFRDIREEEVSNMIDEIRKRSTSSSNSVIDMRDVLS

            CMTNNIINRVTIGRTYNEGESGRAVKALLEDLLALLGTFNIGEYIPWLKWLNKINGLDNR

            VKKVAKDLDAFLDSVIEERVVRNKKAENSTGEAKDFVDVLLEIQNGNETGFPLQRDSLKA

            ILLDSFIAGVDSIYTTLEWIMIELLRNPRAMEKLQNEVRGLVQGKAEITEDDLGNMLYLK

            AVIKESLRLNPPFPIPVPRESMEDVKLLDYNIPAKTQVLVNIWAIGRDPLSWDEPEEYRP

            ERFLNSDIDFRGLNFELIPFGAGRRGCPGIPFAIVIIELTLARLVNKFNFALPQGIKKED

            LDMSECTGISIRRKLPLLAVATPCSV*
```

```
SlCYP736A61     MLYTTPSNLTRRVWFNIESIFSSYSMTLIWTILVVGLSVYALYELLNIHKRKRYPPGPTG
mut             MLYTTPSNNRASHFRTSSFVRQKSTQRFTKSSPKTWSYYVYETRASTYNHCLIC*

                LPILGHLHLLGKNPHKDLQNLAQKHGPIMYMRLGLVPTIIASSADAAEKVLKTYDHIFAS

                RPHHEASQHLAYGQKNLVFAKYDVYWRNIRKLCTVHLLSNQKIHSFQSMRKQEVELLIES

                LKQEARDRVVIDLSAKVTSLNANLTCLMVFGKKYMDEDLDKRGFKAIVQDVVHLAGMPNL

                GDFFPFLGAIDLQGITRKLKDLSKVFDEFLEKIIDEHVYAHEHKQNKDFVDTMMDIMQSG

                DAKFQFDRHHIKAILFDMLIAAMDTTASSTEWILTELLRHPQVMKELQKELQEVVGLDRM

                VEESDLENLKYLDMVVKEGLRLHPVAPLFYHESIEDCVVDGFHIQKGTRIIINCYAIHMD

                PNVWPDPEKFLPERFVGSSVDFRGRDFQLVPFGSGRRSCPGMQLAVMVVRLVVAQLVHCF

                EWELPNGIESCDLDIDEKFGIVTCREKPLLAIPTYRLNK*
```

```
SlCYP733A1      MEAIPGTLGWPIIGESLSFISEFSSPAGIYSFINKRQKLYGKVFKSYVLGRYTVFMTGRE
mut             MEAIPGRYIQLHQQETETVWEGV*

                ASKILLTGKDGIVSLNLFYTGQQVLGPTSLLQQTGEAHKRLRRLIAEPLSVDGLKKYFQF

                INSLAIETLDQWSGRKILVLEETSTFTLKVIGNMIMSLEPTGEEQEKFRTNFKIISGSFA

                SLPFKVPGTAFYRGIQARDRMYAMLDSIIDQRRSGENIKQDFLQSLVKKHGKDAPEGDDD

                DKLTDKQLKDNILTLLVAGHDTTTAALTWLLKFLQENPAVLERLREEHREIQAQKQGTLD

                LTWSEVNNMPYTAKVISETLRMATILPWFSRKAAQDFEIEGCKIKKGWSLNLDVVSIHRD

                PKIFPNPEKFDPSRFDDPLKPFSFLGFGSGPRMCPGINLAKLEISVFLHHLVCRYKWTPL

                DTDDSVQPTLVRMLKNKYPVMVEPL*
```

```
SlUGT37   MSENHPHVLIFPYPAQGHMLPLLDFTHQLVNNGVYITILVTPKNLPFLNPLLSRNPSIKT
mut       MEYTSLF*

          LVLPFPSHPSIPAGVENVKDLPANGFLSMMCNLGKLRDPILEWFHNHPSPPSAIISDMFL

          GFTHEIATQLGIRRYVFSPSGALALSVVYSLWREMPKRKDPNDENENFHFPNIPNSPKFP

          FWQISPIYRSYVEGDPSTEFIRECYLADIASHGIVFNTFIELENVYLDYLMKDLGHNRVW

          SVGPVLPPGEEDDVSVQSNRGGSSSVLASEILAWLDKCEDHSVVYVCFGSQAVLTNKQME

          ELAIALDKSGVHFILSAKRATKGHASNDYGVIPSWFEEKVAGRGLVVRDWAPQVLILKHR

          AIATFLTHCGWNSTLESLTAGIPLLTWPMGADQFANANLLVDEHEVAIRACEGVQTVPNS

          DELAALLAEAVQGNKVEDRRLRASKLRKIAINGIKEGGNSFKELAAFVKHLREEATIIEA

          *
```

pJPC663

```
SlUGT97   MTTHKAHCLILPYPVQGHINPMLQFSKRLRSKRVKITIALTKSFLKNMKELPTSMSIEAI
mut       MTTHKAHCLILPYPVQGHINPMLQFSKRLRSKRVKITIALTKSFLKNMKELPTSMSIEAI

          SDGYDDGGRDQAGTFVAYITRFKEIGSDTLSQLIQKLAISGCPVNCIVYDPFLPWAVEVA
          SDGYDDGGRDSRNFRGLYYTIQRNWFGYSVSTYSKIGN*

          KQFGLISAAFFTQNCVVDNIYYHVHKGVIKLPPTQNDEKIIIPGFSSPIKASDVPNFVIN

          PEGERILEMLVNQFSNLDKVDWVLINSFYELEKEVIDWMSKIYPIKTIGPTIPSMYLDKR

          LHDDKEYGLSMFKPMTNECLNWLNHQPISSVLYVSFGSLAKLGSEQMEELAWGLKNSNKS

          FLWVVRSTEEPKLPNNFIEELTSEKGLVVSWCPQLQVLEHESIGCFLTHCGWNSTLEAIS

          LGVPMVAMPQWSDQPTNAKLVKDVWEIGVRAKQDEKGIVRREVIEECIKLVMEEDKGKLI

          RENAKKWKEIARNVVDEGGSSDKNIEEFVSKLVTIS*
```

SlCYP72A178 MEILYNTIIATICVAILLVYTWRVLNWAWFRPKKLENFLRQRGLKGNPYKLLYGDLNELT
*mut* MEILYNTIIATICVAILLVYTWRVLNWAWFRPKKLENFLRQRGLKGNPYKLLYGDLNELT

KSIVEAKSKSINISDDITQRLIPFFLDSINKNGKSSFMWLGPYPTVLITNPEHVKEILTK
KSIVEAKSKSINISDDITSKAYPFFP*

NYVYLKQTHPNPFAKLLAQGLVLVEEDKWAKHRKIINPAFHVEKLKHMLPAFYMSCSEMI

SKWEDIVSKETSYELDVWPDLQIMTSEVISRTAFGSSYEEGRIVFELQQEQAEHIMDISR

SIYIPGSRFLPTKRNKRMLEIEKQIQTTIRHIIDKRLRAMEAGETSKDDLLGILLESNMK

EIEQHGNKDFGLTTTEVIEECKLFYFAGQETTSVLLVWTMVLLCLHPEWQVRAREEVLQV

FGNEKPDLEGLSHLKIVTMILYETLRLFPPLPVFSRRNKEEVKLGELQLPAEVILIIPAI

FIHYDKEIWGEDAKEFKPERFSEGVSKATKGQVSFIPFGWGPRICIGQNFAMMEAKMAIA

MILQKFSFELSPSYTHAPFATITIHPQYGAPLLLRKH*

SlCYP72A184 MIVTIFCCAIAITLFVCLWKVLNWVWFQPKKLEKLLRKQGLNGNSYRFLYGDMKDFSKMI
*mut* MIVTIFCCAIANYFVCVSMESTKLGLVPTKEVGEIVEETRAKWEFIQIFVWGHERFF*

KEANSKPMNMDDHDLAPRMVPFFLETIKKYGKKCFTWMGPRPQILIMDPELIKEVLSKTY

VYQKPHGNPLGTLFVQGLVSYEKDKWAKHRKIINPAFHLEKLKHMLPAFYLSCSEMLSKW

EDVVRVEGSHEIDVWPHLQQLTCDVISRTAFGSSYEEGRKIFELQKEQAQHFLEVIRSVY

IPGWRFLPTKRNRRMKELKNDVRSSIRGIIDKRLKAMEAGNADNEDLLGILLESNFKEIE

LHGNKDFGMSIEEVIEECKLFYFAGAETTSVWILWTLVLLSRHPDWQERAREEVLQVLGS

RKPDFDGLNHLKVVTMILYESLRLYSPITVLTRRVYEDITLGEVSLPTGVLVSLPMILLH

HDKDIWGEDATKFNPERFSEGISSATKGQVTYFPFAWGPRICIGQNFAILEAKMTLCMIL

QSFSFELSPSYTHAPQSLVTTQPQYGAPLIFHKL*

```
SlCYP72A186   MDEIQILVRVCCSAIAIALFVCLWKVLNWVWLNPKKLENLLRKQGLNGNSYKILYGDLND
mut           MDEIQILVRVCCSAIGSVCVSVESTKLGLAQSEEVGEFVEETRAKWELLQNIVWGFE*

              FIGMIREASSKPMNLSDDIAPRLVPFFLETTKKYGKKCFIWLGPKPQVIIMDPELIKEVL

              SKTYLYQKPGGNPFAALLVQGLATYEEDKWAKHRKIINPAFHLEKLKHMLPAFHLSCTEM

              LSKWEDAVPLGSSREIDVWPHLQQLTCDVISRTAFGSSYEEGRKIFELQTEQAQNFIDAV

              REVYIPGRRFLPTKRNRRMKEIKNEVRTSVKGIIDKRMKAMKAGNADNNEDLLGILLESN

              FKEIEQHGNKDFGMSIEEVIEECKLFYFAGQETTSVLLVWTLVLLSRYQDWQTQAREEVL

              QVFESRKPDFDGLNHLKVVTMILYESLRLYPPLITLNRQVNEDIVLGELCLPAGVLVSLP

              MILLHHDKDIWGEDANEFNPERFREGISSATKGKVTYFPFSWGPRICIGQNFAMLEAKMA

              LCMILQSFSFELSPSYTHAPKSLVTMQPQYGAPLILHKL*
```

#### pJPC723

```
SlUGT111      MASSKFPSIIETLKPNLIIYDGFQPWVATMASSYSIHAIMFYVSSTSGLAYIYHQFLHGS
mut           MASSKFPSIIETLKPNLIIYDGFQPWVATMASSYSIHAIMFYVSSTSGLAYIYHQFLHGS

              SSLTSFPFSSIYLHDHEIKKLGIQPIKPRDEKAFAYIILESFEQSHNIVLLNTCRETEGK
              SSLTSFPFSSIYLHDHEIKKLGIQPIKPRDEKAFAYIILESFEQSHNIVLLNTCRERGSI

              YIDYVSTIGKKELIPIGPLIREAMIDEEEDWGTIQSWLDKKDQLSCVYVSFGSESFLSKQ
              *

              EIEEIAKGLELSKVNFIWTIKFPKGVNKTIEEMVPQGFLESTKSNFEPFKYWRFCNSLWM

              EFDVRKHEFWHTINSHAYES*
```

```
SlUGT112   MASSKFPSIIETLKPNLIIYDGFQPWVATMASSYSIHAIMFYVSSTSGLAYIYHQFLHGS
mut        MASSKFPSIIETLKPNLIIYDGFQPWVATMASSYSIHAIMFYVSSTSGLAYIYHQFLHGS

           SSLTSFPFSSIYLHDHEIKKLGIQPIKPRDEKAFAYIILESFEQSHNIVLLNTCRETEGK
           SSLTSFPFSSIYLHDHEIKKLGIQPIKPRDEKAFAYIILESFEQSHNIVLLNTCRERGSI

           YIDYVSTIGKKELIPIGPLIREAMIDEEEDWGTIQSWLDKKDQLSCVYVSFGSESFLSKQ
           *

           EIEEIAKGLELSKVNFIWTIKFPKGVNKTIEEMVPQGFLESTKGKGMVIEGWAPQSLILN


           HSSIGGFITHCGWNSILESMSFGIPIIAMPMNHDQPLNSRLVEELGIGVEILRGENGKIM


           KEEVAKGIRKVIEEKPRKQIHLKAMQLSEKIKLKAIDEGVKKLLKLLY*
```

```
SlUGT113   MASSKFPSIIETLKPNLIIYDGFQPWVATMASSYSIHAIMFYVSSTSGLAYIYHQFLHGS
mut        MASSKFPSIIETLKPNLIIYDGFQPWVATMASSYSIHAIMFYVSSTSGLAYIYHQFLHGS

           SSLTSFPFSSIYLHDHEIKKLGIQPIKPRDEKAFAYIILESFEQSHNIVLLNTCRETEGK
           SSLTSFPFSSIYLHDHEIKKLGIQPIKPRDEKAFAYIILESFEQSHNIVLLNTCRERGSI

           YIDYVSTIGKKELIPIGPLIREAMIDEEEDWGTIQSWLDKKDQLSCVYVSFGSESFLSKQ
           *

           EIEEIAKGLELSKVNFIWTIKFPKGVNKTIEEMVPQGFLESTKSNFEPFKYWRFCNSLWM


           EFDVRKHEFWHTINSHAYES*
```

```
SlUGT116   MVQPHVLLVTFPAQGHINPSLQFAKRLIEMGIEVTFTTSVFAHRRMAKIAASTAPKGLNL
mut        MVQPHVLLVTFPAQGHINPSLQFAKRLIEMGIEVTFTTSVFAHRRMAKIAASTDTQGLKL

           AAFSDGFDDGFKSNVDDSKRYMSEIRSRGSQTLRDVILKSSDEGRPVTSLVYTLLLPWAA
           GGIL*

           EVARELHIPSALLWIQPATVLDIYYYYFNGYEDEMKCSSSNDPNWSIQLPRLPLLKSQDL


           PSFLVSSSSKDDKYSFALPTFKEQLDTLDGEENPKVLVNTFDALELEPLKAIEKYNLIGI


           GPLIPSSFLGGKDSLESSFGGDLFQKSNDDYMEWLNTKPKSSIVYISFGSLLNLSRNQKE


           EIAKGLIEIQRPFLWVIRDQEEEKEEEKLSCMMELEKQGKIVPWCSQLEVLTHPSLGCFV


           SHCGWNSTLESLSSGVPVVAFPHWTDQGTNAKLIEDVWKTGVRMRVNEDGVVESDEIKRC


           IEIVMDGGEKGEEMRKNAQKWKELARAAVKEGGSSEVNLKAFVLQVSKSC*
```

```
SlUGT129   MKIERKQSVVLVPHPYQGHLTPMLQLGSILHSQGFSVIVAHTQYNTPNYSNHPQFVFHSM
mut        MKIERKQSVVLVPHPRGI*

           DDGLQGIDMSFPSLENIYDMNENCKAPLRNYLVSMMEEEGDQLACIVYDNVMFFVDDVAT

           QLKLPSIVLRTFSAAYLHSMITILQQPEIYLPFEDSQLLDPLPELHPLRFKDVPFPIINN

           TVPEPILDFCRAMSDIGSSVATIWNTMQDLESSMLLRLQEHYKVPFFPIGPVHKMASLVS

           STSILEEDNSCIEWLDRQAPNSVLYVSLGSLVRIDHKELIETAWGLANSDQPFLWVIRPG

           SVSGFQCAEALPDGFEKMVGERGRIVKWAPQKQVLAHPAVAGFFTHCGWNSTLESICEEV

           PMVCRPFLADQLVNARYLSQIYKVGFELEVIERTVIEKTIRKLMLSEEGKDVKKRVADMK

           QKIVAGMQIDCTSHKNLNDLVDFISALPSRLAPPTPVFGAIMSSNHIASKCIIES*


SlUGT135   MGVLTIEPHFVLFPFMAQGHTIPMIDIARLLSQRGVIITIVTTHLNANRFKKVIDRAIES
mut        MGVLTIEPHFVLFPFMAQGHTIPMIDIARLLSQRGVIITIVTTHLNANRFKKVIDRAIES

           GLKIQVVHLYFPSLEAGLPEGCENFDMLPSMDLGLKFFDATERLQPQVEEMLREMKPSPS
           GLKIQVVHLYFPSLEAGLPEGCENFDMLPSMDLGLKFFDATERLQPQVEEMLREMKPSPS

           CIISDMCFPWTTNVAQKFNIPRIVFHGMGCFSLLCLHNLKDWEGLEKIESDTEYFRVPGL
           CIISDMCFPWTTNVAQKFNIPRIVFHGMGCFSLLCLHNLKDWEGLEKIESDTEYFRVPGL

           FDKIELTKNQLGNAARPRNEEWRVMSEKMKKAEEEAYGMVVNTFEDLEKEYIEGLMNAKN
           FDKIELTKNQLGNAARPRNEEWRVMSEKMKKAEEEAYGMVVNTFEDLEKEYIEGLMNAKN

           KKIWTIGPVSLCNKEKQDKAERGNEAAIDEHKCLNWLDSWEQNSVLFVCLGSLSRLSTSQ
           KKIWTIGPVSLCNKEKQDKAERGNEAAIDEHKCLNWLDSWEQNSVLFVCLGSLSFHVSDG

           MVELGLGLESSRRPFIWVVRHMSDEFKNWLVEEDFEERVKGQGLLIRGWAPQVLLLSHPS
           *

           IGAFLTHCGWNSSLEGITAGVAMITWPMFAEQFCNERLIVDVLKTGVRSGIERQVMFGEE

           EKLGTQVSKDDIKKVIEQVMDEEMEGEMRRKRAKELGEKAKRAMEEEGSSHFNLTQLIQD

           VTEQAKILKPM*
```

```
SlCYP94A24  MMIIDLEFSTLLICLVPFLFIFFINLFHSKSTKIPKSYPIIGSYFSILANKHRRIQWSSD
mut         MMIIDLEFSTLLICLVPFLFIFFINLFHSKSTKIPKSYPIIGSYFSILANKHRRI_____

            IIQTTTNLTFTLIRPFGNRQIFTANPSNVQYMLKTHFHIYQKGHMFKSTMADFLGDGIFN
            _______FTFTLIRPFGNRQIFTANPSNVQYMLKTHFHIYQKGHMFKSTMADFLGDGIFN

            VDGEIWKYQRQVASHEFNTKSLRKFIETVVDAELNERLIPLLAVAASEKKVVDLQDILQR
            VDGEIWKYQRQVASHEFNTKSLRKFIETVVDAELNERLIPLLAVAASEKKVVDLQDILQR

            FAFDNICKIAFGYDPAYLSPSLPQEKFAVAFEEAVKLSSERFNAIFPFLWKIKRHFDFGS
            FAFDNICKIAFGYDPAYLSPSLPQEKFAVAFEEAVKLSSERFNAIFPFLWKIKRHFDFGS

            EKRFRIAVDEVRQFAKQLVREKQSELNQKSSLDSVDLLSRFLSVGHRDDDFVTDVVISFI
            EKRFRIAVDEVRQFAKQLVREKQSELNQKSSLDSVDLLSRFLSVGHRDDDFVTDVVISFI

            LAGRDTTSAALTWFFWLISEHPVVESSILNEIKLKSHTPVYEEVKEMIYTHASLCESMRF
            LAGRDTTSAALTWFFWLISEHPVVESSILNEIKLKSHTPVYEEVKEMIYTHASLCESMRF

            YPPVPIDSKTATEDNVLPDGTFVKKGTRVSYHPYAMGRAEELWGSDWNKFKPERWLDKDE
            YPPVPIDSKTATEDNVLPDGTFVKKGTRVSYHPYAMGRAEELWGSDWNKFKPERWLDKDE

            VTGNWNFVTKDMYTYPVFQAGPRICLGKEMAFLQMKRVVAGVLRRFKVVPVTEKGVEPVF
            VTGNWNFVTKDMYTYPVFQAGPRICLGKEMAFLQMKRVVAGVLRRFKVVPVTEKGVEPVF

            ISYLTAKMKGGFHVTIEERDHE*
            ISYLTAKMKGGFHVTIEERDHE*


SlUGT135    MLKTRFDNYPKGKTFSTILGDFLGRGIFNVDGDLWRFQKRMSSLELGKVSIRSYAFEVVK
mut         MLKTRFDNYPKGKTFSTILGDFLGRGIFNVDGDLWRFQKRMSSLELGKVSIRSYAFEVVK

            NEIDKRLVPLLHEYKQGGGVLDFQDVFRRFSFDCICRFSFGLDPKCLESSLPMSQFALSF
            NEIDKRLVPLLHEYKQGGGVLDFQDVFRRFSFDCICRFSFGLDPKCLESSLPMSQFALSF

            DLASKLSAKRAMTTSPIVWKIKRFFNIGSEKELREAIKVINILAQEVIRQKRKLAFSNHR
            DLASKLSAKRAMTTSPIVWKIKRFFNIGSEKELREAIKVINILAQEVIRQKRKLAFSNHR

            DLLSRFMGSISDETYLRDIVISFLLAGRDTIASALTSFFYVIGNNPQVAKAIREEANRVL
            DLLSRFMGSISDETYLRDIVISFLLALRRFHVELAQPYHHNPRFSPGLTASFKAYSTNPN

            GPNEDLTSYEQMSELHYLQACVYESMRLYPPIQFDSKFCLEDDFLLDGTFVKKGTRVTYH
            HGGWNSFQHIN*

            PYAMGRMEELWGCDALEFKPQRWLKDGVFVQENPFKYPVFQAGLRVCLGKEMALVEVKSV


            ALSLLRRFHVELAQPYHHNPRFSPGLTASFKGGLLVSVRQISS*
```

**Figure S2**. Lesions in target genes.

Alignments of predicted protein sequence for wildtype and mutant alleles. Discrepancies highlighted in grey.

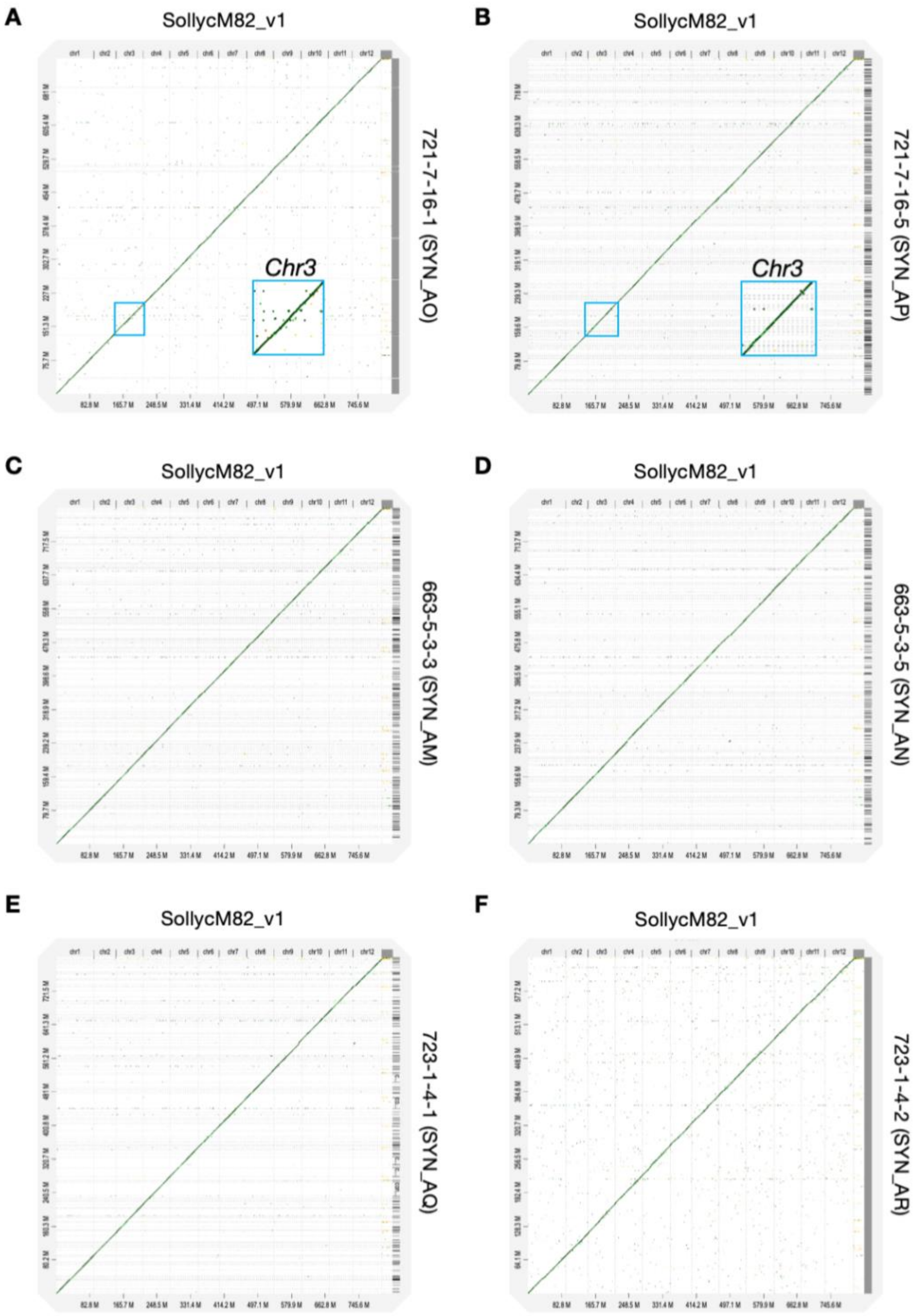


**Figure S3**. Genome assemblies of gene-edited lines.

D-GENIES visualization of mutant genome assemblies mapped to the M82 genome **(A-F)**. Enlarged blue box in **(A)** and **(B)** highlights *Chr3* where an inversion was detected in the 721-7-16 line.

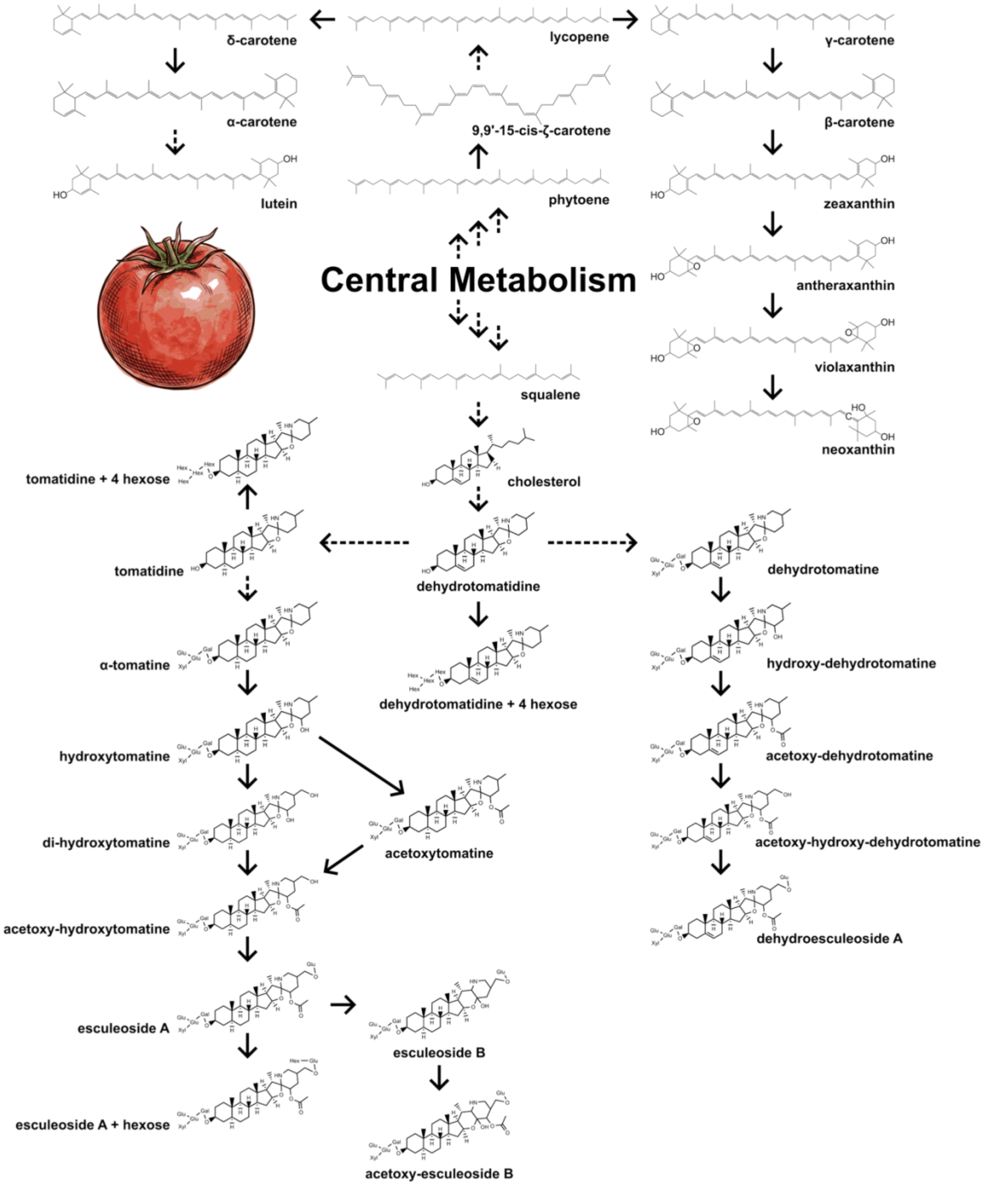


**Figure S4**. SGA and Carotenoid pathways.

Structures of carotenoid and steroidal glycoalkaloids profiled in this study. Solid arrows denote single enzymatic steps.

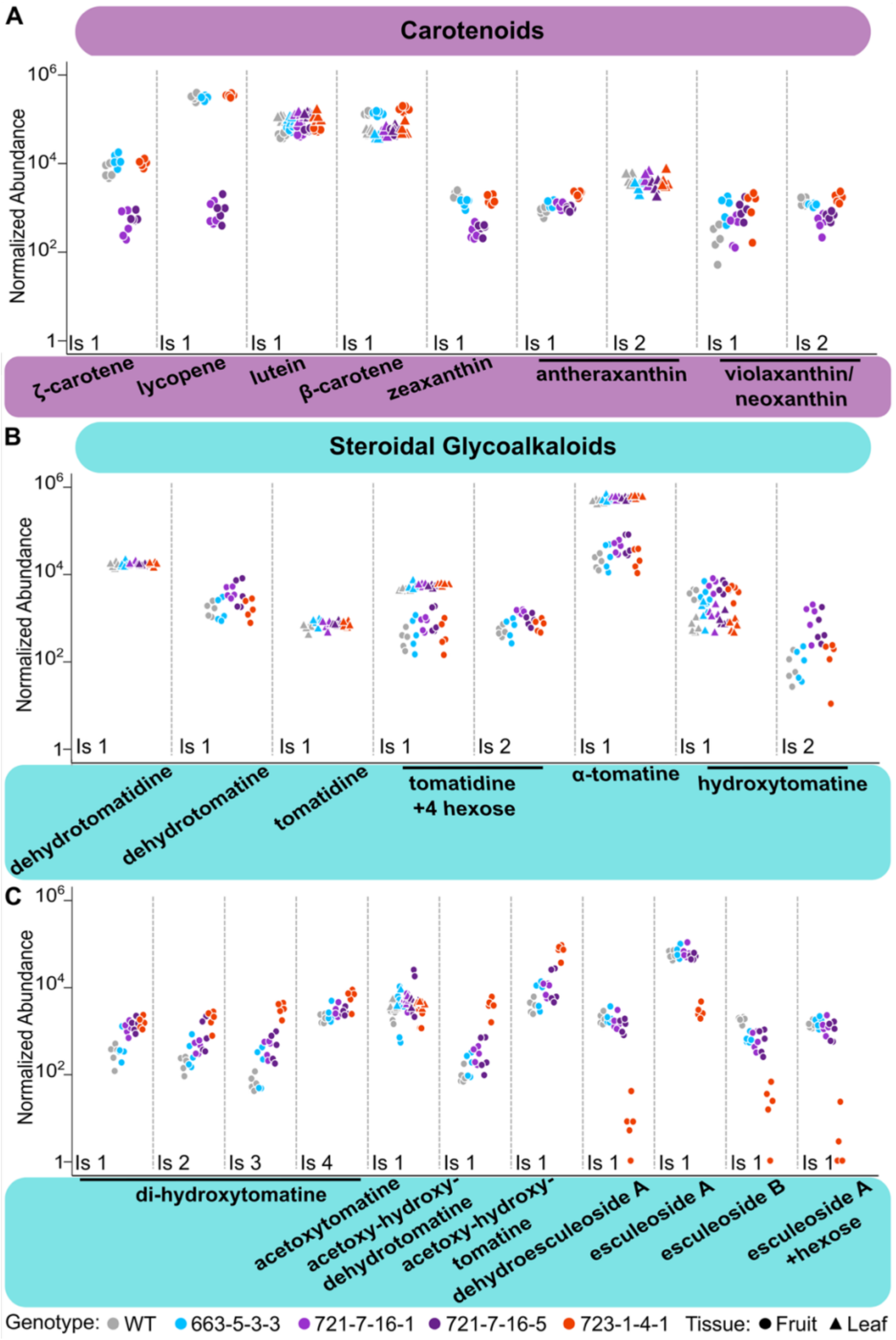


**Figure S5**. Abundance of SGA and carotenoid metabolites in fruit and leaves.

Normalized abundance of **(A)** carotenoid isomers (Is) and **(B)** and **(C)** SGAs in fruit and leaves.
